# Tissue-specific BMAL1 cistromes reveal that enhancer-enhancer interactions regulate rhythmic transcription

**DOI:** 10.1101/319319

**Authors:** Joshua R Beytebiere, Alexandra J Trott, Ben Greenwell, Collin A Osborne, Helene Vitet, Jessica Spence, Seung-Hee Yoo, Zheng Chen, Joseph S Takahashi, Noushin Ghaffari, Jerome S Menet

## Abstract

The mammalian circadian clock relies on the transcription factor CLOCK:BMAL1 to coordinate the rhythmic expression of thousands of genes. Consistent with the various biological functions under clock control, rhythmic gene expression is tissue-specific despite an identical clockwork mechanism in every cell. Here we show that BMAL1 DNA binding is largely tissue-specific, due to differences in chromatin accessibility between tissues and co-binding of tissue-specific transcription factors. Our results also indicate that BMAL1 ability to drive tissue-specific rhythmic transcription not only relies on the activity of BMAL1 cis-regulatory elements (CREs), but also on the activity of neighboring CREs. Characterization of the physical interactions between BMAL1 CREs and other CREs in the mouse liver reveals that interactions are quite stable, and that BMAL1 controls rhythmic transcription by regulating the activity of other CREs. This supports that much of BMAL1 target gene transcription depends on BMAL1 capacity to rhythmically regulate a network of enhancers.

## Introduction

Circadian clocks are found ubiquitously across all kingdoms of life and provide a time-keeping mechanism for organisms to anticipate rhythmic environmental changes. In mammals, virtually every cell harbors the same circadian clockwork that regulates rhythmic gene expression along with temporal cues and systemic signals, such that biological functions occur at the most appropriate time of day. The mammalian circadian clock relies on transcriptional feedback loops initiated by the heterodimeric transcription factor CLOCK:BMAL1 (1, 2). CLOCK:BMAL1 rhythmically binds DNA to drive the rhythmic transcription of the genes *Period* (*Per1, Per2, Per3*) and *Cryptochrome* (*Cry1* and *Cry2*), which upon translation form a repressive complex that feedbacks to inhibit CLOCK:BMAL1-mediated transcription. CLOCK:BMAL1 binds to E-boxes and activates transcription more potently during the middle of the day, and maximal repression occurs during the middle of the night (3, 4). CLOCK:BMAL1 is also responsible for the transcriptional regulation of many clock controlled genes, which allows for rhythms in biochemistry, physiology, and behavior (5).

Characterization of the rhythmic transcriptional outputs driven by the circadian clock indicates that genes expressed in a rhythmic manner vary greatly between tissues in the mouse (6-8). Similar findings in plants, insects, and primates also revealed that circadian gene expression is largely tissue-specific (9-11). However, it is still unknown how the same circadian clock mechanism can generate tissue-specific rhythmic gene expression. Previous work in *Drosophila* revealed that CLOCK:CYC, the CLOCK:BMAL1 homolog, exhibits tissue-specific DNA binding between the body and head, and that much of this tissue specificity is due to the co-operation between CLOCK:CYC and tissue-specific transcription factors (TFs) (9). Tissue-specific binding of another mammalian clock component, *Rev-erba*, has also been described, but the underlying mechanisms have not yet been characterized (12).

To address the mechanisms by which the mammalian circadian clock generates tissue-specific circadian transcriptional programs, we performed BMAL1 chromatin immunoprecipitation at the genome wide level (ChIP-Seq) in the mouse liver, kidney, and heart. Our data revealed that the majority of BMAL1 DNA binding is tissue-specific, and that permissiveness of the chromatin environment and co-binding of tissue-specific TFs (ts-TFs) account for much of this tissue-specificity. In addition, we found large discrepancies between BMAL1 DNA binding and rhythmic gene expression with, for example, many genes targeted by BMAL1 in all three tissues only exhibiting rhythmicity in a single tissue. Characterization of the underlying mechanism suggests that the ability of BMAL1 to drive tissue-specific rhythmic gene expression depends not only on how BMAL1 promotes the activity of the cis-regulatory elements (CREs) it binds to, but also of other neighboring CREs. Together, our data suggest that BMAL1 transcriptional output is controlled through enhancer-enhancer interactions, and that BMAL1-driven rhythmic transcription depends on the capacity of BMAL1 to rhythmically regulate a network of enhancers.

## Results

### BMAL1 DNA binding signal is largely tissue specific

To determine if BMAL1 cistrome differs between tissues, we performed BMAL1 ChIP-Seq in the mouse liver, kidney and heart with three biological replicates per tissue. Because CLOCK:BMAL1 DNA binding sites are virtually identical over the course of the day and only exhibit homogenous differences in DNA binding strength (3, 4), we performed BMAL1 ChIP at the time of maximal binding, *i.e*., ZT6 (Fig. S1A) (ZT stands for Zeitgeber Time, and ZT0 is defined at the light on signal). Identification of BMAL1 DNA binding sites using HOMER software (13) revealed that most BMAL1 peaks were tissue-specific, with less than 6% of BMAL1 peaks common to the liver, kidney and heart (Fig. 1A, B; Table S1). Closer inspection revealed that tissue-specific BMAL1peaks were more often associated with BMAL1 signal below peak detection threshold in the other tissues rather than a complete absence of signal (Fig. 1B; S1B; see below). In addition, quantification of BMAL1 ChIP-Seq signal revealed that stronger BMAL1 binding is associated with peaks common to more than one tissue (Fig. 1C).

**Figure 1.**
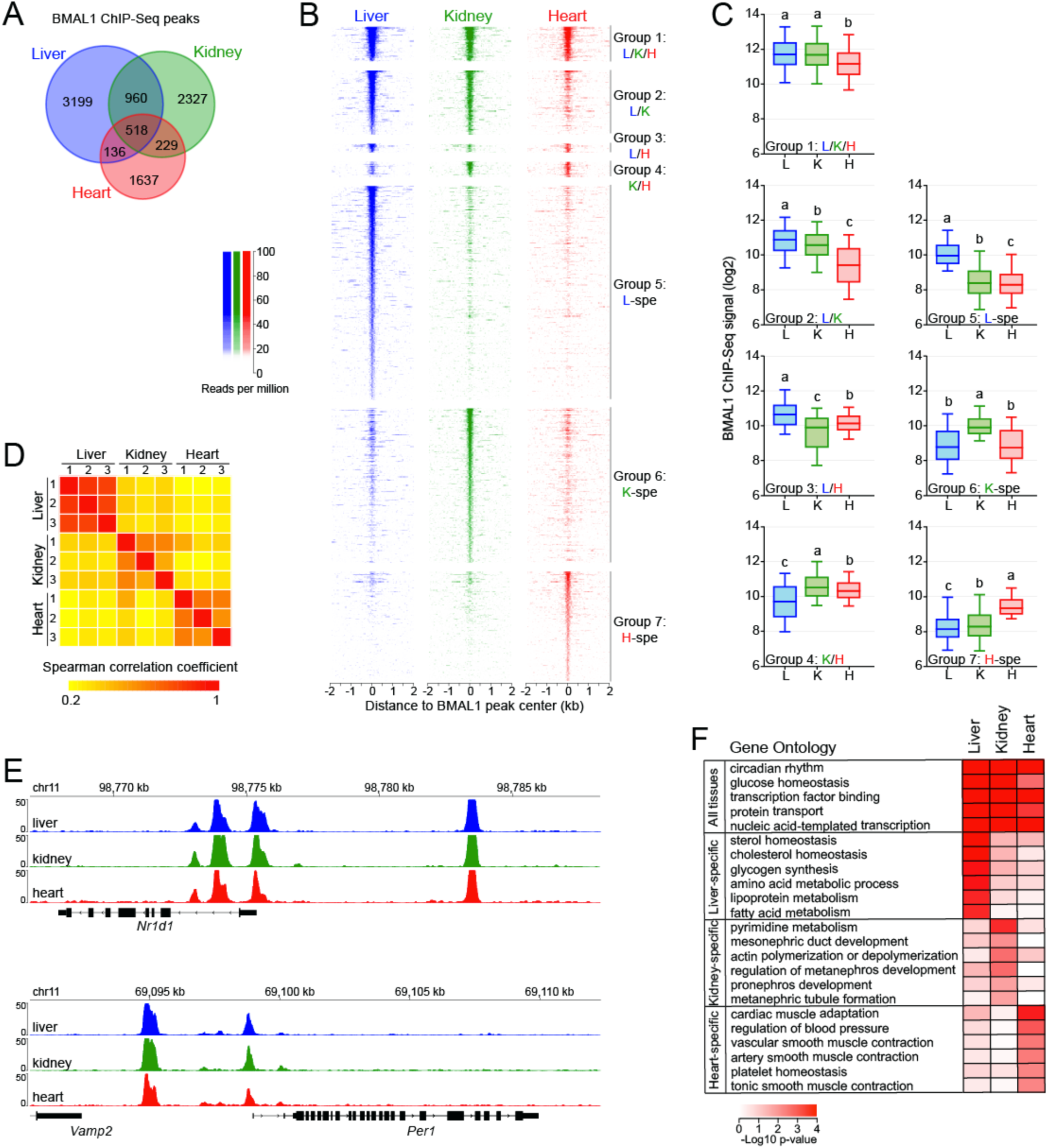
BMAL1 cistromes are largely tissue-specific. **(A)** Overlap of BMAL1 ChIP-Seq peaks in the mouse liver, kidney, and heart. **(B)** BMAL1 ChIP-Seq signal at BMAL1 peak center ± 2kb in the mouse liver, kidney, and heart, and parsed based on tissues in which BMAL1 peaks were detected. Group1: peaks common to all three tissues; group 2: peaks common to the liver and kidney; group 3: peaks common to the liver and heart; group 4: peaks common to the kidney and heart; group 5: liver-specific peaks, group 6: kidney-specific peaks; group 7: heart-specific peaks. **(C)** BMAL1 ChIP-Seq signal calculated at peak center ± 250 bp in the mouse liver, kidney, and heart. Groups with different letters are statistically different (Kruskal-Wallis test ; p < 0.05). **(D)** Spearman correlation coefficients of BMAL1 ChIP-Seq signal between each biological replicate (n = 3 per tissue) calculated at all 9,006 BMAL1 ChIP-Seq peaks. **(E)** Genome browser view of BMAL1 ChIP-Seq signal at *Nr1d1* and *Per1* gene loci. **(F)** Gene ontology analysis performed using BMAL1 peaks detected in the liver, kidney, or heart, and reporting functions enriched across the three tissues or in only one tissue (p-value < 0.05).

To ensure that differences in BMAL1 ChIP-Seq signal between tissues represent true biological variation, we compared BMAL1 ChIP-Seq signal between each of the nine datasets and found a high degree of correlation between samples originating from the same tissue, but not between tissues (Fig. 1D). We also found that BMAL1 ChIP-Seq signal at core clock genes is nearly identical between all three tissues, further indicating that differences in BMAL1 binding signal between tissues stem from biological variations (Fig. 1E). Consistent with this finding, gene ontology analysis of BMAL1 target genes for each tissue revealed that while circadian rhythm and other general terms are enriched in all three tissues, tissue-specific biological processes are enriched in a tissue-specific manner (Fig. 1F). Together, these results indicate that BMAL1 predominantly binds DNA in a tissue-specific manner, and targets genes involved in the regulation of tissue-specific biological functions.

### Chromatin accessibility contributes to tissue-specific BMAL1 DNA binding

To understand the large differences in BMAL1 ChIP-Seq signal between tissues, we set out to define the mechanisms that enable BMAL1 to bind DNA in a tissue-specific manner. Characterization of the chromatin landscape by the ENCODE consortium and others revealed that 94.4% of transcription factor (TF) ChIP-Seq peaks are located in an accessible chromatin region, *i.e*., a region that is hypersensitive to DNase I (14, 15). Because mouse liver CLOCK:BMAL1 DNA binding sites are predominantly located in DNase I hypersensitive sites (DHS) (16, 17), we hypothesized that tissue-specific BMAL1 binding may be, at least in part, caused by differences in DNA accessibility between the liver, kidney and heart. To test our hypothesis, we used DNase-Seq datasets from the mouse ENCODE project (15) and found that many tissue-specific BMAL1 peaks are located in a chromatin region that is only accessible in the corresponding tissue (Fig. 2A). Quantification of DNase-Seq signal at liver-, kidney-, and heart-specific BMAL1 peaks (named group 5, 6, and 7 hereafter, respectively) confirmed that tissue-specific BMAL1 peaks are associated with higher DNase-Seq signal (Fig. 2B). However, large differences in DNase-Seq signal suggested that tissue-specific BMAL1 peaks may not be always associated with a complete absence of DNase-Seq signal in other tissues.

**Figure 2.**
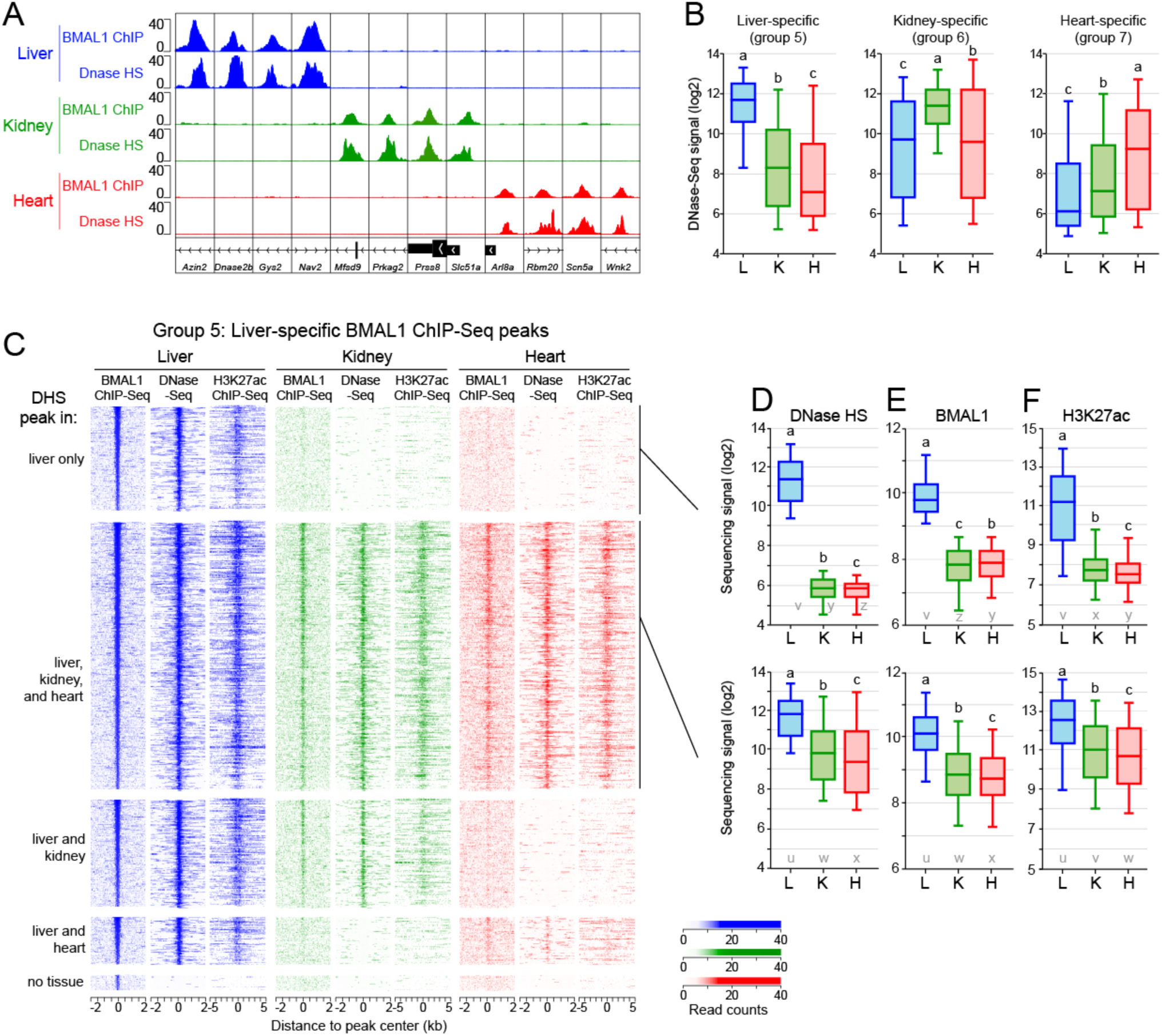
The chromatin environment shapes tissue-specific BMAL1 binding. **(A)** Genome browser view of BMAL1 ChIP-seq and DNase-seq signals in the mouse liver, kidney, and heart at twelve BMAL1 tissue specific peaks. **(B)** DNase-seq signal calculated at BMAL1 peak center ± 250 bp in the mouse liver, kidney, and heart for tissue specific BMAL1 peaks. Groups with different letters are statistically different (Kruskal-Wallis test; p < 0.05). **(C)** BMAL1 ChIP-seq, DNase-seq, and H3K27ac ChIP-seq signal at liver specific BMAL1 peaks, parsed based on the presence of a DNase I hypersensitive sites (DHS) peak in the liver, kidney, and heart. BMAL1 ChIP-seq and DNase-seq signals are displayed with a window of ± 2kb, whereas H3K27ac ChIP-seq signal is displayed with a window of ± 5kb. **(D-F)** Quantification of DNase-seq (D), BMAL1 ChIP-Seq (E), and H3K27ac ChIP-Seq (F) signals for liver specific BMAL1 peaks located at (top, group 5A) liver specific DHS or (bottom, group 5B) DHS peaks common to the liver, kidney and heart. Groups with different letters are statistically different (Kruskal-Wallis test; p < 0.05). Letters u-z denotes the outcome of the statistical analysis performed using groups 5A and 5B together.

To investigate if tissue-specific BMAL1 binding occurs at DHS common to several tissues, we mapped the genomic location of DHS in the liver, kidney and heart (Fig. S2A), and determined the fraction of tissue-specific BMAL1 peaks found at tissue-specific DHS or at DHS common to more than one tissue, focusing primarily on liver-specific BMAL1 peaks. As expected, almost all mouse liver BMAL1 peaks (97.7%) are located in chromatin regions that are accessible in the mouse liver (Fig. 2C). However, only ~20% of liver-specific BMAL1 peaks are located at a liver-specific DHS, *i.e*., most liver-specific BMAL1 peaks are located within chromatin regions that are also accessible in other tissues (Fig. 2C). Importantly, we found that while no BMAL1 signal is detectable in the kidney and heart at liver-specific DHS, some BMAL1 signal is visible at common DHS even if it is not defined as significant BMAL1 peaks by HOMER (Fig. 2C-E). Similar results were observed for kidney- and heart-specific BMAL1 peaks (Fig. S2B, C).

Because DNase I hypersensitivity identifies accessible chromatin regions independently of their transcriptional activity, we also analyzed public ChIP-Seq datasets for histone modifications found at transcriptionally active/primed cis-regulatory elements (CREs), *i.e*, acetylated lysine 27 of histone 3 (H3K27ac) and monomethylated lysine 4 of histone 3 (H3K4me1). Remarkably, both H3K27ac and H3K4me1 ChIP-Seq signals mirrored the DNase-Seq signal, and were in particular detected at kidney and heart DHS exhibiting liver-specific BMAL1 peaks (Fig. 2C, 2F, S2D).

In summary, our data indicate that only a minority of BMAL1 binding sites are located at tissue-specific DHS. For most tissue-specific BMAL1 peaks, chromatin is accessible in other tissues and harbors histone modifications that mark transcriptionally active CREs, albeit to lower levels. Importantly, some BMAL1 DNA binding can be detected at those sites, although at levels too low to be categorized as ChIP-Seq peaks by peak-calling algorithms.

### Tissue-specific BMAL1 peaks are enriched for tissue-specific transcription factor DNA binding motifs

Several mechanisms may contribute to tissue-specific BMAL1 binding. First, BMAL1 DNA binding may only rely on the presence of its binding motif in an accessible CRE. Under this scenario, BMAL1 binding signal would correlate with DNase-Seq signal, and differences between tissues would mostly depend on the proportion of cells harboring an accessible CRE and/or the extent to which the CRE is accessible. Alternatively, tissue-specific BMAL1 binding may be, as suggested for some other TFs (9, 18, 19), facilitated by tissue-specific TFs (ts-TFs) that bind at sites nearby BMAL1. Here, BMAL1 DNA binding signal would not correlate well with DNase-Seq signal, and motifs for ts-TFs would be enriched at tissue-specific BMAL1 peaks.

To assess if either or both of these mechanisms promote tissue-specific BMAL1 binding, we first compared BMAL1 ChIP-Seq signal with DNase-Seq signal at liver-, kidney-, and heart-specific BMAL1 peaks, and found that both signals were correlated in the heart, but not in the liver and kidney (Fig. 3A). Differences in BMAL1 ChIP-Seq signal of more than 16-fold are detected at liver DHS displaying similar DNase-Seq signals, and similar BMAL1 ChIP-Seq signals were observed at CREs harboring differences of DNase-Seq signal of more than 30-fold in both liver and kidney (Fig. 3A). Importantly, these differences are independent of the number of E-boxes within each CRE (Fig. 3B, S3A). These results thus indicate that, at least for the liver and kidney, the extent to which the chromatin is accessible cannot solely explain the differences in tissue-specific BMAL1 binding.

**Figure 3.**
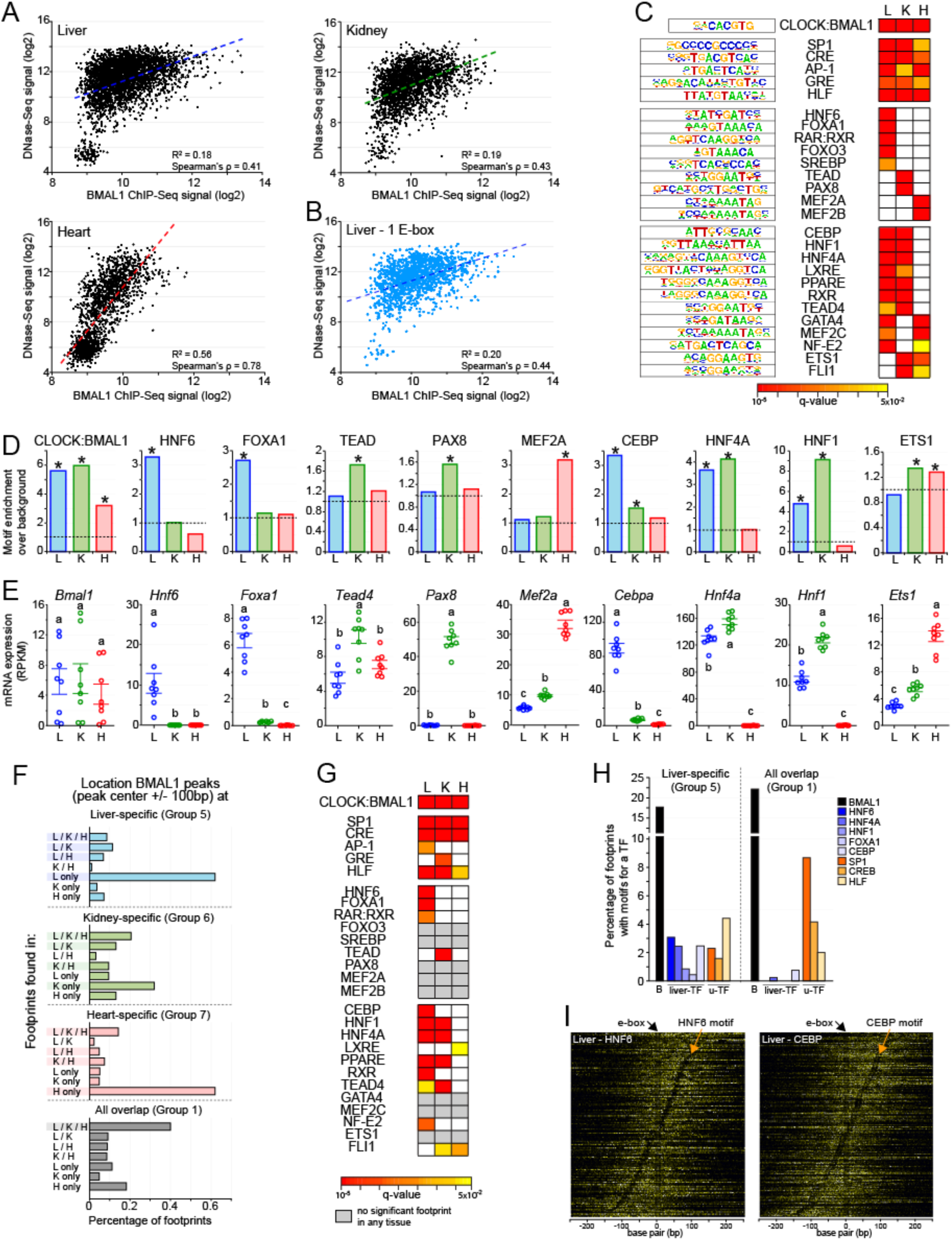
Motifs and footprints for tissue-specific transcription factors are enriched at tissue-specific BMAL1 enhancers. **(A)** Correlation between DNase-seq and BMAL1 ChIP-seq signals (calculated at BMAL1 peak center ± 250 bp) at liver-, kidney-, and heart-specific BMAL1 peaks. **(B)** Correlation between DNase-seq and BMAL1 ChIP-seq signal for liver specific BMAL1 peaks harboring one E-box only. **(C)** Enrichment of TF motifs at liver-, kidney-, and heart-specific BMAL1 peaks, performed using HOMER at BMAL1 peak center ± 100 bp. Enrichments are shown if q-value < 0.05. **(D)** Fold-enrichment of TF motif over background at tissue-specific BMAL1 peaks (BMAL1 peak center ± 100 bp; q-value <0.05). **(E)** mRNA expression in the mouse liver, kidney, and heart of *Bmal1* and TFs whose motifs were enriched at BMAL1 ChIP-Seq peaks. (RNA-Seq datasets from 8). Groups with different letters are significantly different (one-way ANOVA; p-value <0.05). **(F)** Distribution of DNase I footprints identified using pyDNase and detected at BMAL1 peak center ± 100 bp, and parsed based on the tissue(s) they were found in. **(G)** Motif enrichment of TFs performed at DNase I footprints identified in liver-, kidney-, and heart-specific BMAL1 peaks (footprint center ± 15 bp). Enrichments are displayed if q-value < 0.05, and colored in grey if no significant footprint is detected in any of the three tissues. **(H)** Percentage of footprints for BMAL1 (black), liver-specific TFs (blue), and u-TFs (orange) identified at liver-specific BMAL1 peaks (group 5) or at BMAL1 peaks common to all three tissues (group 1). **(I)** DNase I cuts at BMAL1 peaks containing an E-box and an HNF6 motif (left) or a CEBP (right). DNase I cut signal is centered on the E-box and sorted based on the distance between the E-box and the liver-specific TF motif.

To determine if ts-TFs may be involved, we performed a motif enrichment analysis at tissue-specific BMAL1 peaks using HOMER. As expected, E-boxes are enriched at tissue-specific BMAL1 peaks for all three tissues, along with a few additional motifs for ubiquitously expressed TFs (u-TFs) (Fig. 3C, 3D, Table S2). However, the majority of the motifs were significantly enriched in only one or two tissues, with many of them corresponding to binding sites of well-known ts-TFs like *Foxa1* and *Mef2a/b* (Fig 3C, 3D). To verify that these TFs are legitimate ts-TFs, we analyzed their levels of expression in the liver, kidney and heart using public mouse RNA-Seq datasets (8) and the Genotype-Tissue Expression (GTEx) portal for human tissues (20). For almost all of the >20 TFs we examined, mRNA levels were specific to one or two tissues, confirming that these TFs are true ts-TFs (Fig 3E, S3B, S3C). Importantly, the tissue-specific pattern of expression matched particularly well with the tissue-specificity of the motif enrichment, suggesting that these ts-TFs bind DNA in a tissue-specific manner and contribute to tissue-specific BMAL1 DNA binding.

### Genomic footprints for tissue-specific transcription factor are specifically enriched at tissue-specific BMAL1 peaks

The concordance between ts-TF motif enrichment and expression pattern (Fig. 3C-E, S3C) does not directly show that ts-TFs bind at BMAL1 DHS in a tissue-specific manner. To further investigate this possibility, we took advantage of the genomic footprints left by chromatin-associated proteins following treatment with DNase I (21). This analysis, which has been used successfully to define the genome-wide DNA binding profiles of dozens of TFs (22-24), exploits DNase-Seq datasets to uncover regions within CREs that are protected from DNase I activity by proteins bound to DNA (Fig. S4A, S4B). To this end, we analyzed the mouse liver, kidney, and heart DNase-Seq datasets using pyDNase (25) and identified the genome-wide location of TF footprints.

This analysis revealed that most footprints found at tissue-specific BMAL1 peaks are significantly enriched in the corresponding tissue. For example, 60% of the footprints found at liver-specific and heart-specific BMAL1 peaks are only detected in the liver and heart, respectively (Fig. 3F, S4C). Conversely, a large fraction of the footprints identified at common BMAL1 peaks are detected in all three tissues (Fig. 3F, S4C). To identify the proteins generating these footprints, we performed a motif analysis restricted to the footprint genomic locations found at liver-, kidney- and heart-specific BMAL1 peaks. While motifs for CLOCK:BMAL1 and some u-TFs are enriched in all three tissues, most significantly enriched motifs correspond to ts-TFs binding sites (Fig. 3G, S4D, and Table S3). For example, footprints are enriched at motifs for *Hnf6, Foxa1, Cebp, Hnf1*, and *Hnf4a* in the mouse liver, and for *Hnf1, Hnf4a*, and *Tead* in the kidney. We noticed that several ts-TFs whose motifs are enriched at BMAL1 DHS (Fig. 3C) do not exhibit a significant footprint (*e.g., Foxo3, Srebp*, Pax8, Gata4, and Ets1; Fig. 3G), suggesting that they do not bind to BMAL1 DHS despite of the presence of a cognate motif. However, the absence of footprints may also reflect the inability of these TFs to generate distinctive footprints because of DNA residence time and DNase I cut bias, as previously shown for a few other TFs (26, 27).

To further assess if ts-TFs may contribute to tissue-specific BMAL1 binding, we determined the relative enrichment of footprints for several ts-TFs and u-TFs at BMAL1 peaks either specific to the liver (group 5) or common to all three tissues (group 1). We found that footprints for ts-TFs were almost always detected at liver-specific BMAL1 peaks, but not at common BMAL1 peaks (Fig. 3H, S4E). Conversely, footprints for u-TFs were enriched at BMAL1 peaks common to all three tissues, when compared to liver-, kidney-, or heart-specific BMAL1 peaks (Fig. 3H; Fig. S4E). Differences in footprint enrichment between ts-TFs and u-TFs were also observed at liver-specific BMAL1 peaks based on whether DHS are liver-specific or ubiquitous, respectively (Fig. S4F). Finally, visualization of DNase I cuts with heatmaps revealed that most BMAL1-bound DHS exhibit footprints for E-box and ts-TFs, suggesting that ts-TFs are bound to DNA concurrently with CLOCK:BMAL1 (Fig. 3I, S4G).

Together, our findings indicate that ts-TFs are more effectively recruited to tissue-specific BMAL1 peaks whereas u-TFs are more effectively recruited to BMAL1 peaks common to different tissues, suggesting that they both contribute to CLOCK:BMAL1 DNA binding.

### BMAL1 peaks common to all three tissues exhibit stronger BMAL1 signal, harbor more E-boxes, and are enriched at promoters

While the majority of BMAL1 binding sites are tissue-specific, more than 500 BMAL1 ChIP-Seq peaks targeting 294 genes are common to the liver, kidney and heart (Fig. 1A). These genes comprise all expected core clock genes and regulate biological processes that are ubiquitous to most mammalian tissues (Fig. S5A).

Consistent with the hypothesis that BMAL1 binding at peaks common to several tissues is less dependent on ts-TFs than tissue-specific BMAL1 peaks, we found that BMAL1 ChIP-Seq signal at common BMAL1 peaks is well correlated with DNase-Seq signal (Fig. S5B), and between tissues (Fig. 4A). Analysis of the DNA binding motifs at common BMAL1 peaks revealed that E-boxes were enriched along with motifs for the clock genes *Rev-erbα/β* and *Rorα/β/γ* (RORE) and *E4bp4/Dbp/Tef/Hlf* (D-box), as well as motifs for well-characterized u-TFs such as CREB Response Element, SP1 and HLF (Fig. 4B, 4C). Importantly, enrichment for E-boxes at common BMAL1 peaks is higher than at tissue-specific peaks (Fig. 3C), possibly contributing to the increased BMAL1 ChIP-Seq signal at those sites (Fig. 1C). To investigate this possibility further, we quantified BMAL1 ChIP-Seq signal based on the number of E-boxes and found that BMAL1 signal was positively correlated with increasing number of E-boxes for all three tissues (Fig. 4D). Moreover, dual E-boxes, *i.e*. two E-box motifs separated by 6 or 7 nucleotides and which have been proposed to be a preferred CLOCK:BMAL1 binding site (28), are also more enriched at common BMAL1 peaks (Fig. 4E). Finally, characterization of the location of BMAL1 peaks revealed that peaks common to all three tissues were predominantly located at promoter/TSS, whereas tissue-specific BMAL1 peaks were more consistently located within introns and intergenic regions (Fig. 4F).

**Figure 4.**
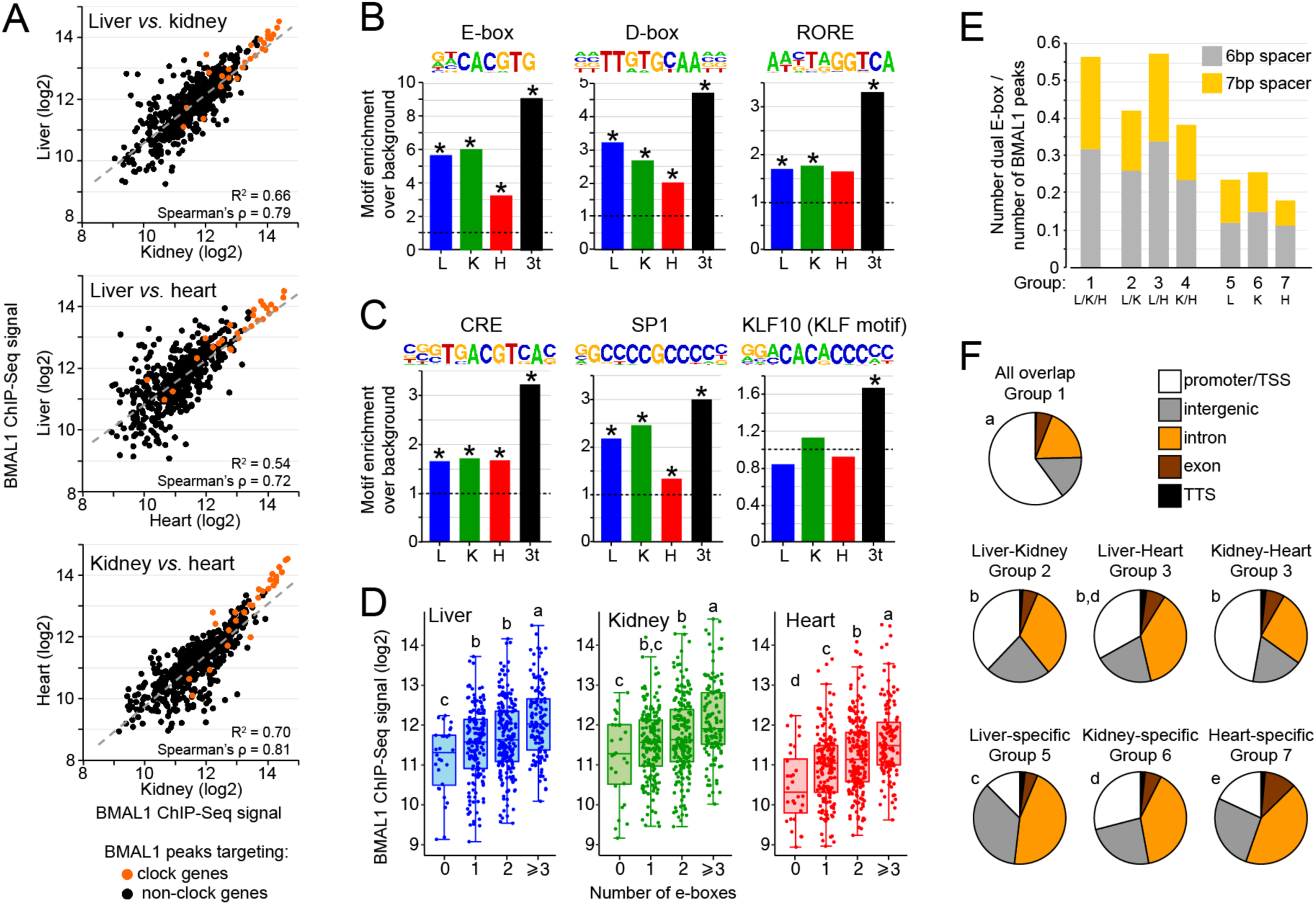
BMAL1 DNA binding sites common to the mouse liver, kidney, and heart exhibit unique features. **(A)** Correlation of BMAL1 ChIP-seq signal between the mouse liver, kidney, and heart for BMAL1 peaks common to all three tissues (group 1). **(B, C)** Enrichment for the motifs of clock genes (B) and u-TFs (C) at tissue-specific BMAL1 peaks (liver in blue, kidney in green, and heart in red) or BMAL1 peaks common to the three tissues (black). Asterisks illustrate a q-value < 0.05. **(D)** BMAL1 ChIP-Seq signal for BMAL1 peaks common to all tissues parsed based on the number of E-boxes (canonical E-box CACGTG and degenerated E-boxes CACGTT and CACGNG). **(E)** Fraction of BMAL1 peaks harboring a dual E-box motif (two E-boxes separated with 6 or 7 bp). **(F)** Location of BMAL1 peaks for each of the seven groups. Statistical analysis was performed using a chi square test, and *post-hoc* analysis was performed with Fisher’s exact test. Groups with different letters are statistically different (p < 0.01).

Together, these results indicate that BMAL1 peaks common to all three tissues, which target genes that are involved in generic biological processes, are enriched for E-boxes and other clock gene motifs, and are predominantly localized at promoter regions/TSS. These features likely contribute to the higher BMAL1 ChIP-Seq signal observed at those sites.

### BMAL1 DNA binding contributes only partially to rhythmic gene expression

To determine whether tissue-specific BMAL1 binding contributes to generating tissue-specific circadian transcription, we analyzed BMAL1 target gene expression in the mouse liver, kidney and heart using public datasets (8). In agreement with the literature (4, 29), genes targeted by BMAL1 exhibit a higher proportion of rhythmic expression than those not targeted by BMAL1. Specifically, rhythmic expression of BMAL1 targets in the liver, kidney and heart is increased by ~2-fold compared to all genes, and by ~1.3-1.5-fold compared to genes targeted by BMAL1 in other tissues (Fig. 5A). Surprisingly, genes targeted by BMAL1 in all three tissues (group 1) do not exhibit substantially more rhythmic expression despite increased BMAL1 ChIP-Seq signal, increased number of E-boxes, and preferential peak location at promoter/TSS (Fig. 5A). Moreover, the majority of BMAL1 target genes are not rhythmically expressed in any of the three tissues (Fig. 5A). These results thus suggest that although BMAL1 DNA binding contribute to increased rhythmic expression, it is not sufficient to drive rhythmic transcription.

**Figure 5:**
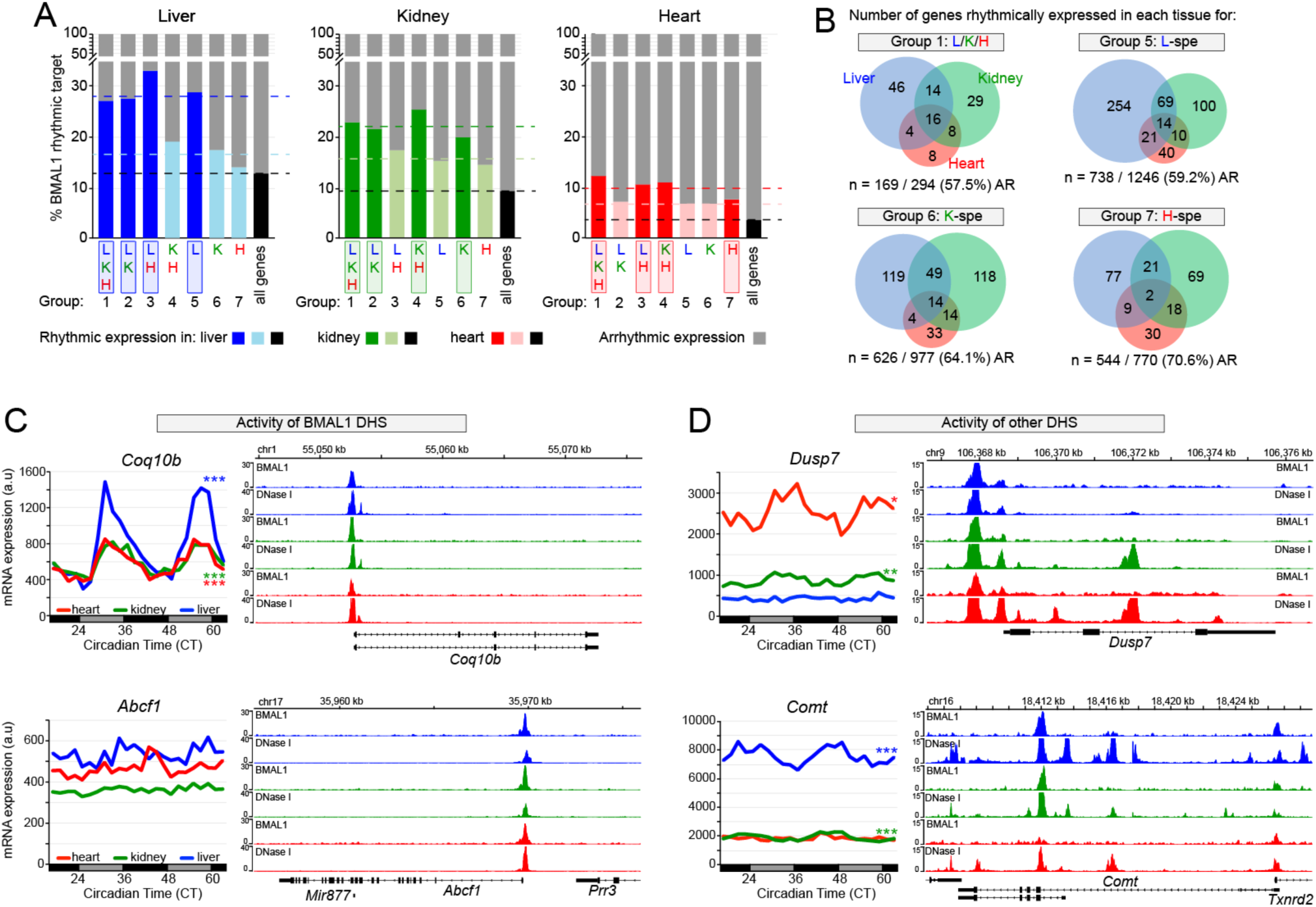
Transcriptional activities of BMAL1 DHS and other DHS contribute to BMAL1-mediated rhythmic transcription. **(A)** Percentage of rhythmically expressed gene for the seven groups of BMAL1 binding sites. Gene expression data originate from public microarray datasets (8) and is considered rhythmic if q-value <0.05 using JTK-cycle. **(B)** Overlap of rhythmic expression for peaks common to all three tissues (group1), and for liver-, kidney-, and heart-specific BMAL1 peaks (groups 5, 6, and 7, respectively). The number of genes that are rhythmically expressed for the mouse liver (blue), kidney (green), and heart (red) is displayed, along with the number and percentage of non-rhythmically expressed genes (AR). **(C, D)** mRNA expression (left) and genome browser view (right) of BMAL1 ChIP-seq and DNase-seq signals in the mouse liver (blue), kidney (green), and heart (red). Rhythmic expression determined by JTK-cycle is defined as *** if q-value < 0.001, ** if q-value < 0.01, and * if q-value < 0.05. The genes *Coq10b* and *Abcf1* (C) represent two examples for which the activity of BMAL1 DHS likely contributes to the differences in mRNA expression, where the genes *Dusp7* and *Comt* (D) represent two examples for which the activity of other DHS likely contributes to BMAL1-mediated rhythmic transcription.

Consistent with this notion, BMAL1 targets common to all three tissues display a widely heterogeneous rhythmic output, with most genes being rhythmic in only one tissue (83 genes out of 125 rhythmically expressed genes; Fig. 5B). In fact, only 16 genes targeted by common BMAL1 peaks are rhythmically expressed in all three tissues, and most of them include clock genes. We also found that many genes targeted by BMAL1 in one tissue were rhythmic only in another tissue (Fig. 5B). To characterize the mechanisms underlying this poor overlap between BMAL1 binding and rhythmic gene expression, we examined the profiles of DNase-Seq and BMAL1 ChIP-Seq datasets in the liver, kidney, and heart. While we were unable to detect a unified mechanism that may explain the variability in BMAL1 transcriptional output, we identified two important features that likely contribute to rhythmic transcription. First, similar BMAL1 ChIP-Seq signal between tissues can result in significantly different levels and/or rhythmicity of target gene expression (Fig. 5C, S6A). This observation was made at genes targeted by a unique DHS, suggesting that factors other than just BMAL1 contribute to the transcriptional activity of BMAL1-bound CRE and to rhythmic BMAL1 target gene expression. Second, we found that rhythmicity of BMAL1 target genes was often associated with higher levels of expression and more DHSs, thus suggesting that BMAL1 CRE(s) functionally interacts with other CRE(s) to regulate transcription activation and drive oscillations in gene expression (Fig. 5D, S6B). We also found a few instances where more DHSs were associated with decreased expression, suggesting that interactions between multiple CREs may also result in decreased transcription activation (Fig. S6C).

In summary, our results indicate that, as recently reported (17), BMAL1 DNA binding is not sufficient to drive rhythmic transcription. They also suggest that rhythmic BMAL1 target gene expression likely results from (i) interactions between CREs bound by BMAL1 and other CREs, and (ii) the transcriptional activity of each CRE, including those bound by BMAL1.

### BMAL1-bound CREs physically interact with other CREs in a constitutive manner

To determine the extent to which BMAL1 target gene transcription relies on physical interactions between CREs, we performed RNA Polymerase II (Pol II) Chromatin Interaction Analysis by Paired-End Tag Sequencing (ChIA-PET) in the mouse liver at ZT6 and ZT18 with 3 biological replicates per time point. By incorporating a Pol II ChIP step, this technique identifies interactions between enhancers and/or TSS of genes that are being transcribed (Fig. 6A) (30, 31).

**Figure 6:**
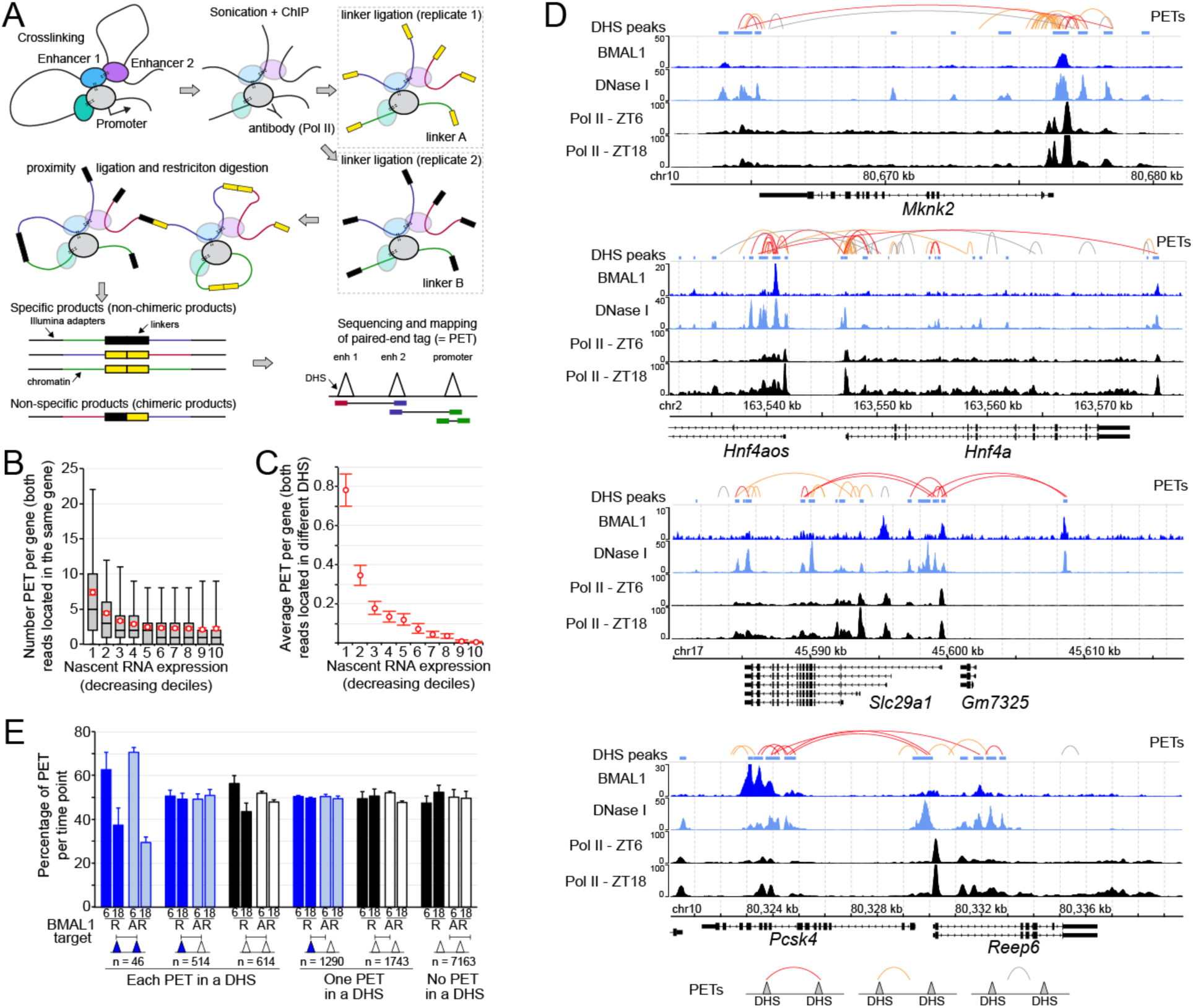
Interactions between BMAL1-bound DHS and other DHS are not rhythmic over the course of the day. **(A)** Illustration of the ChIA-PET technique. **(B)** Number of Paired-End Tags (PET) with both reads mapped to the same gene and parsed based on gene nascent RNA expression in the mouse liver. Red circles represent the average PET number for each decile ± 95% confidence intervals. **(C)** Average number of PET per gene in the mouse liver with both reads located in different DHS and mapped to the same gene, parsed based on gene nascent RNA expression. Error bars represent the 95% confidence intervals. **(D)** Genome browser view of mouse liver BMAL1 ChIP-seq (this study), DNase-seq (ENCODE), and Pol II ChIP-seq (from 16). PET with both reads mapped to DHS are in red, while PETs with one read mapped to a DHS are in orange and those not mapped to a DHS are in grey. **(E)** Percentage of PET detected at ZT6 or ZT18 (average ± s.e.m. of 3 independent experiments), displayed based on the class of PET. Triangle filled in blue represents a DHS with a BMAL1 ChIP-Seq peak, while triangle filled in white represents a DHS without a BMAL1 peak.

To assess the efficiency of our mouse liver Pol II ChIA-PET in detecting chromatin interactions at transcriptionally active genes, we determined the number of Paired-End Tag (PET) mapped to a single gene based on gene transcription using public mouse liver Nascent-Seq datasets (29). We found that most PET mapped to transcriptionally active genes, with 23.4% of all PET located within the top 10% of genes transcribed in the mouse liver (Fig. 6B, S7A, Table S4). Analysis of PET with reads located in two different DHS and visualization of chromatin interactions at mouse liver BMAL1 target genes further confirmed that actively transcribed genes harbor many PET mapping to different DHS, including some bound by BMAL1 (Fig. 6C, 6D). These data indicate that CREs located within the same gene physically interact together via gene looping, and suggest that Pol II transcriptional activity results from the synergistic interaction between these CREs.

A few recent reports suggested that rhythmic interactions between BMAL1 CRE and other CREs underlie the rhythmic transcription of BMAL1 target genes (32-34). To investigate this possibility, we quantified the number of PET based on time of mouse liver collection, the presence of a BMAL1 ChIP-Seq peak, and rhythmic gene expression. Surprisingly, we found that the number of PET at ZT6 is not different than at ZT18, and this even for PET mapped to BMAL1-bound DHS and targeting a gene that is rhythmically transcribed (Fig. 6E). Interactions between two BMAL1-bound DHS were more enriched at ZT6 than ZT18; though the number of PET for this specific category of BMAL1 target genes was low (Fig. 6E).

Our analysis confirms that transcriptionally active genes exhibit physical contacts between CREs. These physical interactions do not appear to be as dynamic as previously reported (32-34), since no significant differences were observed between the time of maximal vs. minimal BMAL1 DNA binding. While this may reflect differences in the techniques used (*i.e*., Pol II ChIA-PET *vs*. 4C -Circularized Chromosome Conformation Capture) and/or level of analysis (e.g., genome-wide *vs*. a few genes), this may also reflect some more fundamental mechanisms into how CREs synergize to regulate the transcriptional activity of their target genes (see discussion).

### Functional interactions between BMAL1-bound CRE and other CRE is associated with rhythmic gene expression

Based on our results suggesting that BMAL1-bound CREs interact with neighboring CREs to drive rhythmic transcription (Fig. 5, 6), we hypothesized that BMAL1 regulates the activity of other CREs to drive rhythmic transcription. To test this hypothesis, we used public mouse liver H3K27ac ChIP-Seq datasets performed in wild-type and *Bmal1^-/-^* mice over the course of the day (16) and examined H3K27ac signal at PET mapped to a BMAL1-bound CRE.

We first focused our analysis on PET having only one tag in a BMAL1-bound CRE and found that, in wild-type mice, H3K27ac signal at BMAL1 CREs is rhythmic with a phase matching BMAL1 DNA binding when target genes are rhythmically expressed (Fig. 7A). Signal at those sites is decreased and arrhythmic in *Bmal1^-/-^* mice, indicating that CLOCK:BMAL1 significantly contributes to H3K27ac rhythmic signal (Fig. 7A). Remarkably, H3K27ac signal at interacting CREs mirrors the signal observed at BMAL1 CREs: it is rhythmic in wild-type mice, and decreased and arrhythmic in *Bmal1^-/-^* mice (Fig. 7A). This therefore suggests that, at rhythmically expressed BMAL1 targets, CLOCK:BMAL1 contributes to the transcriptional activity of the CREs it physically interacts with. Importantly, these results are not observed at arrhythmically expressed BMAL1 targets. H3K27ac signal does not fully match BMAL1 rhythmic DNA binding profile (*e.g*., lowest H3K27ac signal at ZT10), and is much less affected by *Bmal1* knockout (Fig. 7A). Similar observations were made at PET having both tags lying in a BMAL1-bound DHS (Fig. 7B). At CREs targeting rhythmically expressed genes, H3K27ac signal is rhythmic in wild-type mice and significantly decreased and arrhythmic in *Bmal1^-/-^* mice. Conversely, at CREs targeting arrhythmically expressed genes, H3K27ac signal is almost constant throughout the day in wild-type mice, and levels are not impaired by *Bmal1^-/-^* knockout (Fig. 7B). Together, these results support a model whereby BMAL1-mediated rhythmic transcription relies on the capacity of BMAL1-bound CREs to functionally regulate the neighboring CREs it physically interacts with.

**Figure 7:**
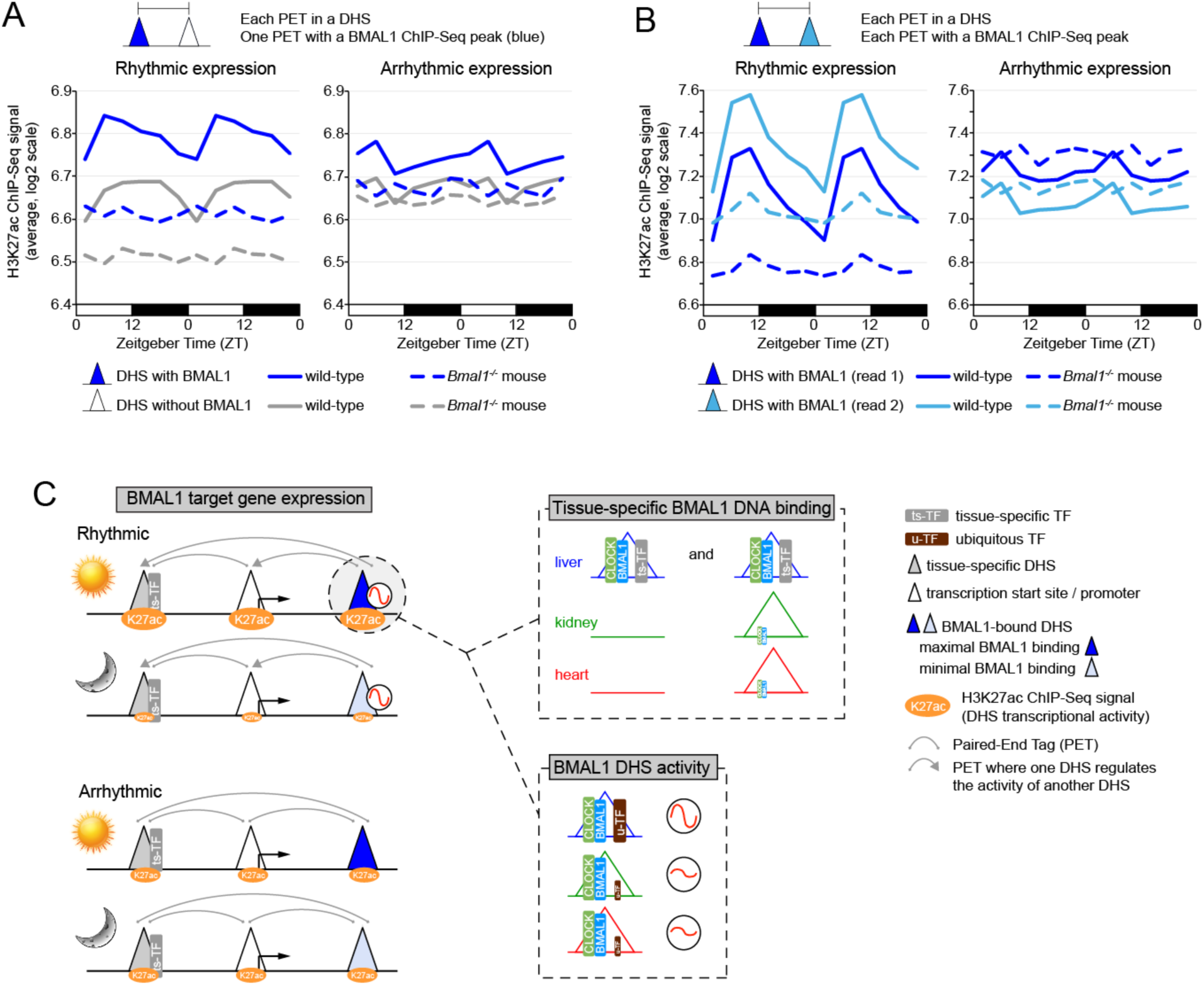
Functional interactions between BMAL1-bound CRE and other CRE is associated with rhythmic gene expression. **(A, B)** H3K27ac ChIP-Seq signal in the mouse liver of wild-type and *Bmal1^-/-^* mice (16). Analysis was performed at PET where both reads are located in a DHS, and is displayed for PET where only one DHS is bound by BMAL1 (A), or for PET where both DHS are bound by BMAL1 (B). Signal was calculated at DHS peak center ± 1 kb, log2-transformed, quantile normalized, and averaged for each group and time points. Distinction is made between BMAL1 target genes being rhythmically expressed (left), or not (right). The 24-hr profile of H3K27ac signal is double-plotted to improve visualization. **(C)** Hypothetical model illustrating how CLOCK:BMAL1 generate tissue-specific rhythmic transcriptional programs. This model incorporates mechanisms on how BMAL1 binds to DNA in a tissue-specific manner (chromatin accessibility and co-binding with ts-TFs), and how u-TFs might contribute to BMAL1 DHS rhythmic transcriptional activity. It also illustrates how functional interaction between BMAL1 DHS and other DHS (including tissue-specific DHS) may contribute to rhythmic transcription.

## Discussion

Although the same clockwork mechanism is found in essentially all tissues, clock-controlled gene rhythmic expression is mainly tissue-specific, reflecting the large number of biological functions that are clock controlled (6-8, 11). By characterizing BMAL1 cistromes in three mammalian tissues, we show that tissue-specific circadian transcriptional programs can be directly initiated by the core circadian clock. We provide evidence that BMAL1 DNA binding profiles are largely tissue-specific, which coincides with tissue-specific chromatin accessibility and co-binding with ts-TFs. Our analysis of BMAL1 target gene expression also indicates that rhythmic CLOCK:BMAL1 transcriptional output relies, at least in part, on the interplay between BMAL1-bound CREs and other CREs. Given that some of these other CREs are tissue-specific, our data suggest that enhancer-enhancer interactions can drive rhythmic BMAL1 target gene expression in a tissue-specific fashion. Taken together, chromatin accessibility and enhancer-enhancer interactions may explain how the circadian clock generates tissue-specific circadian transcriptional programs, thereby regulating biological functions in a tissue-specific manner. We anticipate that these mechanisms apply to other TFs that regulate gene expression in a tissue-specific manner.

Recent characterization of the chromatin accessible landscape revealed that most TFs bind to regions that are hypersensitive to nuclease digestion (14, 15). Because many of these DHS are tissue-specific (Fig. S2A), it is not surprising that differences in the openness of the chromatin between tissues can promote tissue-specific BMAL1 binding. However, the majority of tissue-specific BMAL1 peaks are located in chromatin regions accessible in other tissues (Fig. 2C, S2C, S2D). CREs are clusters of TF binding motifs, and TF recruitment to CREs often results from cooperative binding between TFs. In particular, ts-TFs can cooperate with u-TFs to facilitate DNA binding: CEBPB cooperates with glucocorticoid receptor in the mouse liver (35), FOXA1 and GATA3 with the estrogen receptor in primary breasts tumors (36, 37), and OPA and SERPENT with *Drosophila* CLK:CYC (9). Our finding that binding motifs and genomic footprints for liver-specific TFs are enriched at liver-specific BMAL1 peaks, but not at common peaks, strongly suggests that this ts-TF/u-TF cooperation applies to BMAL1 and that ts-TFs facilitate tissue-specific CLOCK:BMAL1 binding (Fig. 7C).

Genes targeted by BMAL1 in all three tissues do not exhibit a higher rate of rhythmic expression and display a remarkable heterogeneity in the rhythmic output (Fig. 5A, 5B), indicating that even robust BMAL1 binding is not sufficient to drive rhythmic transcription. As recently suggested (17), this may be explained by the contribution of u-TFs in regulating the transcriptional activity of BMAL1 CREs and promoting gene expression. However, our analysis suggest an additional level of regulation that includes CREs unbound by BMAL1, and through which interactions between BMAL1-bound CREs and other CREs would contribute to the regulation of rhythmic gene expression (Fig. 5D).

To characterize how CREs not bound by BMAL1 might contribute to rhythmic BMAL1 transcriptional output, we conducted mouse liver Pol II ChIA-PET to identify physical interactions between DHS within genes engaged in active transcription (Fig. 6). Surprisingly, the number of interactions was found to be similar between the time of maximal and minimal BMAL1 binding. These results are in contrast with previous reports that used the 4C technique and showed rhythmic interactions between BMAL1 enhancers (32-34). However, this discrepancy may be due to differences between the ChIA-PET and 4C techniques or in the number of BMAL1 CREs that were analyzed (e.g., genome-wide *vs*. a few genes). It is also possible that the 4C results preferentially identified interactions between two BMAL1-bound CREs, for which we also observed rhythmic interactions (Fig. 6E). In any case, functional analysis of CRE activity using H3K27ac signal revealed that, at rhythmically expressed target genes, BMAL1 significantly contributes to the transcriptional activity of the CREs it binds to, but also of the CREs it interacts with (Fig. 7A). In particular, we found a striking decrease of H3K27ac signal in *Bmal1-/-* mice not only at BMAL1-bound CREs, but also at interacting CREs (Fig. 7A). Given that the effects of BMAL1 on CRE transcriptional activity were limited at arrhythmically expressed genes, our results support the conclusion that rhythmic gene expression is associated with BMAL1 ability to regulate the activity of its enhancers as well as other enhancers.

Based on these results, we propose a model whereby BMAL1-mediated transcription relies on functional interactions between BMAL1-bound CREs and other CREs (Fig. 7C). These physical interactions would be stable over the course the day for most BMAL1 target genes, thereby enabling CLOCK:BMAL1 to regulate the activity of other enhancers throughout the 24-hr day (*i.e*., not just at time of maximal CLOCK:BMAL1 DNA binding), and preventing these other CREs from driving clock-independent transcription during times of CLOCK:BMAL1 repression. This model is consistent with the expression of CLOCK:BMAL1 target genes in *Bmal1^-/-^* mice which, for most genes, is arrhythmic and accumulates at intermediate levels (29, 38). Indeed, the lack of BMAL1 in *Bmal1^-/-^* mice likely decreases the impact of CLOCK:BMAL1-bound DHS, thus relieving other CREs from the regulation of BMAL1-bound CREs and leading to intermediate levels of gene expression. We also propose that rhythmic transcription depends on the ability of CLOCK:BMAL1 to generate a strong CRE that can coordinate and/or override the transcriptional activity of the other interacting CREs. Failure to produce strong circadian enhancers would generate BMAL1 enhancers that exhibit arrhythmic transcriptional activity. Such a model can provide a mechanistic framework that might explain, at least in part, how changes in the environmental conditions (e.g., high-fat diet and antibiotics treatment in the liver, LPS treatment in the lung; 39, 40, 41) can reprogram circadian transcriptional programs. For instance, environmental challenges may reshape the strength of specific CREs by either increasing or decreasing the transcriptional capabilities of some TFs, thereby altering the functional interactions between CREs and rhythmic gene expression. Future experiments will be required to test these possibilities.

In summary, our results provide novel insights into how BMAL1 regulates rhythmic gene expression in a tissue-specific manner, and shed light on the role of enhancer-enhancer interactions in generating circadian transcriptional programs. We anticipate that these findings will be relevant for our understanding of how the circadian clock regulates a wide array of biological functions under normal and diseased states, and will apply more generally to how other TFs regulate tissue-specific gene expression.

## Materials and methods

### Animals

Male C57BL/6J and *Bmal1-/-* mice were housed under 12-h light:12-h dark (LD12:12) with food and water available *ad libitum. Bmal1-/-* were kindly provided by Christopher Bradfield (42). The age of animals collected was between three and six months old. All experiments were approved by the TAMU Institutional Animal Care and Use Committee (AUP# 2016-0199 and AUP 2013-0158).

### BMAL1 chromatin immunoprecipitation

Mice were euthanized by isoflurane anesthesia followed by decapitation, and livers, kidneys, and hearts were collected, rinsed in ice-cold 1X PBS, minced, and immediately homogenized in 1X PBS containing 1% formaldehyde for 10 minutes at room temperature (4ml for livers, 2ml for kidneys, and 1ml for heart). Formaldehyde cross-linking was quenched by adding 2M glycine to a final concentration of 140 mM. Samples were then kept on ice for ten minutes, washed twice with hypotonic buffer (10mM Hepes pH 7.6, 15mM KCl, 0.15% NP-40, 1mM DTT, and 1mM PMSF), and centrifuged at 1500g for 2 minutes at 4°C. Nuclei were purified by centrifuging on a sucrose cushion (10mM Hepes pH7.6, 15mM KCl, 0.15% NP-40, 24% sucrose, 1mM DTT, 1mM PMSF) at 20,000g for 10 minutes at 4°C, and then washed with hypotonic buffer four times. Sonication for the liver and kidneys were done by resuspending the samples in 12ml per liver and 4.5 ml per kidney of sonication buffer (10mM Tris pH 7.5, 150mM NaCl, 2mM EDTA, 0.25% SDS, 0.2% Triton). Sonication of the heart was done by resuspending the heart in 500 µl of sonication solution (10mM Tris pH 7.5, 150mM NaCl, 2mM EDTA, 0.5% Sarkosyl, 1X protease inhibitor cocktail). Samples were sonicated in 500 µl aliquots to obtain chromatin fragments of about 100-600 bp in length. After sonication, samples were centrifuged at 15,000g for 10 minutes at 4°C, supernatants were moved to a new tube, and inputs (25 µl) and ChIP (200 µl) samples were made. The 200 µl ChIP aliquots were diluted (final concentration: 10mM Tris-Cl pH 7.5, 150mM NaCl, 1% Triton X-100, 0.1% sodium deoxycholate, 0.1% SDS or sarkosyl, 2mM EDTA), 1 µl of BMAL1 antibody (chicken anti-BMAL1) was added and left to rotate overnight at 4°C. Dynabeads antibody coupling kit (#14311D, Invitrogen) was used with rabbit anti-chicken IgY antibody (# 31104, Invitrogen) to immunoprecipitate BMAL1 chromatin complexes. Dynabeads were washed with IP buffer twice (10mM Tris-Cl pH 7.5, 150mM NaCl, 1% Triton X-100, 0.1% sodium deoxycholate, 0.1% SDS or sarkosyl, 2mM EDTA), resuspended in blocking solution (IP buffer with 1 mg/ml bovine serum albumin and 0.1 mg/ml yeast tRNA) and left rotating overnight at 4°C. After overnight incubation, dynabeads were washed once with final IP buffer, the chromatin was added, and left rotating at 4°C for two hours. BMAL1 immunoprecipitated chromatin was then washed twice with TSEI buffer (10mM Tris pH 7.5, 0.1%SDS, 1% Triton X-100, 2mM EDTA, 150mM NaCl, 1mM DTT, 1X Protease Inhibitor Cocktail), twice with TSEII buffer (10mM Tris pH 7.5, 0.1%SDS, 1% Triton X-100, 2mM EDTA, 500mM NaCl, 1mM DTT, 1mM PMSF), twice with LiCl Buffer III (10mM Tris pH 7.5, 0.25M LiCl, 1% NP-40, 1% Na Deoxycholate, 1mM EDTA, 1mM DTT, 1mM PMSF), twice with TENT buffer (10mM Tris pH 7.5, 1mM EDTA, 150NaCl, 0.1% Triton X-100), and once with TET buffer (10mM Tris pH 7.5, 1mM EDTA, 0.1% Triton). ChIP samples were then resuspended in 200uL of ChIP Elution Buffer (50mM Tris-HCl pH8, 10mM EDTA, 1% SDS, 1mMDTT) while input samples were supplemented with 175 µl of ChIP Elution buffer, and incubated for 6-18 hours at 65°C. After the incubation, samples were purified using Qiagen PCR purification columns (# 28106), and efficiency of BMAL1 ChIP was verified by qPCR as described below.

### Generation of BMAL1 ChIP-seq libraries and sequencing

BMAL1 and input ChIP-Seq libraries were generated from liver, kidney, and heart samples (n = 3 mice per tissue) using NEBNext^®^ ChIP-Seq Library Prep Master Mix Set (# E6240, NEB) as per the manufacturer’s instructions. DNA from ChIP and input were quantified using a Quantus Fluorometer (# E6150, Promega), and 10 ng was used to generate the libraries. DNA end repair was performed with NEBNext End Repair Reaction Buffer and Enzyme Mix for 30 minutes at 20°C. dA-Tailing of end-repaired DNA was performed with NEBNext dA-Tailing Reaction Buffer and Klenow Fragment (3’→5’ exo) for 30 minutes at 37°C. Adapter ligation of dA-tailed DNA was performed with Quick Ligation Reaction Buffer, NEBNext Adaptor (1.5 µM), and Quick T4 DNA Ligase. Libraries were generated by PCR amplification of adaptor ligated DNA using NEBNext Multiplex oligonucleotides and Phusion Taq (M0530S). Libraries were amplified for 16 cycles. Libraries were quantified with qPCR with TRUseq library standards, and with a Quantus Fluorometer. DNA cleanup between each reaction was performed using Solid Phase Reversible Immobilization (SPRI) beads generated in the lab from Sera-mag SpeedBeads (# catalog number 09-981-123, Thermo-Fisher). BMAL1 ChIP-seq and input libraries were sequenced using an Illumina NextSeq with a sequence length of 76 bp.

### BMAL1 ChIP-qPCR

Chromatin Immunoprecipitation was performed as described above with the following exceptions. After tissue collection, tissues were flash frozen in liquid nitrogen and kept at -80°C. Nuclei extraction was conducted using seven samples at a time (6 time points in wild-type mice -ZT2, ZT6, ZT10, ZT14, ZT18, and ZT22-, and the ZT6 sample from *Bmal1-/-* mouse) to minimize inter-individual variations, and nuclei were flash-frozen in glycerol storage buffer (10mM Tris-Cl pH 7.5, 50mM NaCl, 2mM EDTA, 50% glycerol, 1mMDTT, 0.15mM spermine, 0.5mM spermidine, 1X PIC). BMAL1 ChIPs were also performed using seven samples at a time as described above, except for the BMAL1 antibody (# ab3350, abcam) and the Dynabeads protein G (# 10004D, Invitrogen).

qPCR was performed using the BIO-RAD iTaq Universal SYBR Green Supermix (# 1725124, and BIO-RAD CFX Connect). ChIP fold enrichment was calculated as the ratio between *Dbp* 1^st^ intron ChIP signal normalized to input and intergenic region ChIP signal normalized to input. The primer sequences used are:

*Dbp* 1^st^ intron (forward): ATGCTCACACGGTGCAGACA
*Dbp* 1^st^ intron (reverse): CTGCTCAGGCACATTCCTCAT
Intergenic region (forward): CTTTTAATGAGGCTGTGTGGA
Intergenic region (reverse): ACTCCCTTGCGAATGTCCTA

### Sequencing datasets and alignment to the mouse genome

All public datasets were downloaded from NCBI or https://encodeproject.org (unless noted below) as fastq or Short Read Archive (SRA) file formats. To avoid issues due to the utilization of different protocols/procedures for each tissue, each of the liver, kidney and heart ChIP-Seq, DNase-Seq and mRNA expression datasets were generated for the from the same research laboratory. Accession number are as follow:

- H3K27ac ChIP-seq (comparison between tissues): downloaded from https://encodeproject.org, accession numbers_GSM1000093 (heart), GSM1000140 (liver), and GSM1000092(kidney).
- H3K4me1 ChIP-seq: downloaded from https://encodeproject.org, accession numbers GSM769025 (heart), GSM769023 (kidney), and GSM769015 (liver).
- DNase-seq: downloaded from https://encodeproject.org, accession numbers GSM1014166 (heart), GSM1014193 (kidney) and (liver).
- RNA-seq and microarray datasets: downloaded from the NCBI website, accession numbers GSE54652.
- H3K27ac and RNA Polymerase II ChIP-seq (time-course in wild-type and *Bmal1^-/-^* mice in the mouse liver): downloaded from the NCBI website, accession number GSE60430.

SRA files were converted to fastq files using SRA toolkit (43). BMAL1 ChIP-seq, DNase-seq, H3K27ac ChIP-seq, H3K4me1 ChIP-seq and Pol II ChIP-Seq were aligned to the mouse genome (version mm10) using bowtie2 (44) using the parameters: -x and --end-to-end. Uniquely mapped reads were only considered for analysis, and up to 3 duplicated sequences were kept for each BMAL1 ChIP-Seq dataset. The heart BMAL1 ChIP-Seq replicate 1 dataset contains two technical replicates that were merged into a single bam file, which was then randomly downsampled using samtools from 47,936,219 read to 17,253,305 reads to avoid overrepresentation of one replicate in the final merged file comprising the three biological replicates. All bam files from the heart were then merged for a total read count of 36,975,623 reads (replicate 1: 17,253,305 reads; replicate 2: 11,994,180 reads; replicate 3: 7,728,138 reads). The kidney biological replicate 1 was also downsampled using samtools from 26,840,611 reads to 8,856,833 reads. All bam files from the kidney were then merged for a total of 26, 930,789 reads (replicate 1: 8,856,833 reads; replicate 2: 7,381,236 reads; replicate 3: 10,692,720 reads). None of the liver samples were downsampled, and the bam files were merged using samtools for a total of 34,846,537 reads (replicate 1: 10,966,215 reads; replicate 2: 11,265,139 reads; replicate 3: 12,615,183 reads). Visualization files were generated using bedtools (45) and normalized to 10,000,000 reads. Input files were processed individually and then merged as bam files using samtools.

For DNase-seq datasets, bam files from all technical and biological replicates were merged and no downsampling was performed. Reads from the RNA-Seq dataset were trimmed using fastx_trimmer (http://hannonlab.cshl.edu/fastx_toolkit/) with the following parameters -f 1 -l 100 -Q 33, and aligned to the genome using STAR (46) with the default parameters and the option: --outFilterIntronMotifs RemoveNoncanonica. Gene expression data were retrieved using cufflinks (47) and the genome version GRCm38.p5_M14 and default parameters.

### Sequencing datasets analysis

#### BMAL1 ChIP-seq and DNase-seq peak calling

Peak calling for both BMAL1 ChIP-seq and DNase-seq data was performed with findPeaks from the HOMER suite (13). A minimum local enrichment of 4-fold was set up as necessary, and the following parameters were used: -style factor and -i (for BMAL1 ChIP-Seq) or -style dnase and -region (for DNase-Seq). No input was used to identify DNase-seq peaks. Overlap between ChIP-Seq or DNase-Seq peaks was determined using the function intersectBed from Bedtools suite (45) using default parameters. Heatmaps were using the Rscript pheatmap.R.

#### Assignment of BMAL1 peaks to their target genes

BMAL1 peaks were assigned to their target genes using the perl script annotatePeaks.pl from the HOMER suite with the mm10 mouse genome as a reference (13). The HOMER gene annotation script outputs each peak into the following categories: (i) Promoter-TSS, corresponding to TSS - 10kb to TSS + 1kb, (ii) Transcription termination site (TTS), corresponding to TTS - 100 bp to TTS + 1kb, (iii) exons, (iv) introns, and (v) intergenic, which corresponds to peaks located upstream of the TSS by more than 10 kb and downstream the TTS by more than 1 kb. Intron and exon assignments were not adjusted from the output of HOMER annotatepeaks.pl. BMAL1 peaks labeled as intergenic were not assigned to a target gene.

#### Quantification of ChIP-Seq and DNase-Seq signal at BMAL1 ChIP-Seq peaks

ChIP-seq and DNase-seq signal was calculated using scripts from Bedtools (45), the uniquely mapped reads (mm10 version) and the genomic coordinates of BMAL1 ChIP-seq peaks (Table S1) or mouse liver DHS peaks (Table S5). Signal was calculated in a ± 250 bp region from the peak center for BMAL1 ChIP-seq signal and DNase-seq signal, and in a ± 1 kb region from the peak center for H3K27ac ChIP-Seq signal. For the analysis of H3K27ac ChIP-Seq signal in wild-type and *Bmal1^-/-^* mouse liver over the course of the day, signal retrieved at DHS peak center ± 1 kb for all 67,433 DHS peaks was quantile normalized for all datasets.

#### Footprint detection

Detection of footprints was performed using the python script wellington_footprints.py from the pyDNase suite (25, 48). All parameters were set to default, and a p-value of -20 was used along with an FDR of 0.01. Wig files generated by wellington_footprints.py were converted to BigWig files with the wigToBigWig algorithm downloaded at https://genome.ucsc.edu/.

#### Motif analysis at BMAL1 peaks and footprints at BMAL1 peaks and enhancers

Motif analysis was performed at BMAL1 DNA binding sites (genomic location of BMAL1 ChIP-Seq peaks) using the perl script findMotifsGenome.pl from the HOMER suite (13), using the parameter -size 200. Motifs were considered as significantly enriched if the q-value was less than 0.05. Motif enrichment was calculated based on the background from the output of findMotifsGenome.pl. Motif analysis at footprints located with BMAL1 peaks and BMAL1 DNase I hypersensitive sites (peak center ± 15bp) has been performed similarly.

#### Detection of E-box and dual E-box motifs

Generation of motifs for E-boxes and dual E-boxes was performed with the perl script seq2profile.pl from the HOMER suite (13). The E-boxes considered for analysis were as follows CACGTG, CACGNG and CACGTT. These E-boxes contained only one mismatch from the canonical motif and were found to be functional CLOCK-binding motifs *in vitro* (49). The dual E-box motif tolerates up to two mismatches between the two E-boxes and contains a spacer of either six or seven base pair.

#### Gene ontology analysis

Gene ontology was performed using the perl script annotatePeaks.pl from the HOMER suite (13), with the parameter -go and mm10 genome. HOMER assigns target genes based on the closest gene to the BMAL1 binding region, and then searches for enriched functional categories.

#### Rhythmic expression analysis

Rhythmic expression of BMAL1 target genes was determined using public microarray datasets performed in the same research lab (8). Files containing expression values for each microarray probe were downloaded from the NCBI website (GSE54652), and no analysis of the original files were performed. Rhythmic gene expression was determined using JTK cycle (ref) with the following parameters: timepoints 18-64, and all other parameters were left to default, and was considered significant if q-value < 0.05. Genes targeted by two or more BMAL1 ChIP-Seq peaks assigned to different categories (*e.g*., a gene targeted by two BMAL1 peaks: one common to all three tissues, and one specific to the liver) were not considered for this analysis.

#### Data from GTEX portal

Graphs in Fig. S3C displaying the RPKM values of different TFs in human liver, kidney, and heart (atrial appendage and left ventricle) were retrieved from GTEX portal (20) in April 2016.

### Mouse liver Pol II ChIA-PET

Pol II ChIA-PET experiments were performed using mouse livers collected at either ZT6 or ZT18, with three biological replicates per time point and following previously published protocols (50-52). Pol II ChIP, which is the starting point of the ChIA-PET experiment (Fig. 6A), was performed similarly to BMAL1 ChIP with the following modifications:

i. Livers were collected, rinsed in ice-cold 1X PBS, flash frozen in liquid nitrogen, and stored at -80°C. Frozen livers were crushed to a fine powder in liquid nitrogen using a mortar, and resuspended in 1X PBS containing the crosslinking reagents. One ChIA-PET experiment (JM11) was performed on livers crosslinked with 1% formaldehyde for 10 minutes at room temperature (single crosslinking), and two experiments (JM08 and JM12) were performed on livers first crosslinked with 1.5 mM EGS for 20 minutes at room temperature, and then for 10 additional minutes with 1% formaldehyde (dual crosslinking).
ii. Nuclei were sonicated in 10 mM Tris pH 7.5, 150 mM NaCl, 2 mM EDTA, 0.5% SDS, 0.2% Triton, 1X protease inhibitor cocktail. Because large amounts of starting material were required, sonicated liver chromatin from 3-7 mice were pulled together, resulting in chromatin amounts ranging from 1.3 mg to 4.1 mg in a volume of ~10 ml. Chromatin samples were diluted 5-fold to obtain a final concentration of ChIP buffer of 10mM Tris-Cl pH 7.5, 150mM NaCl, 1% Triton X-100, 0.1% sodium deoxycholate, 0.1% SDS, 1X protease inhibitor cocktail. Pol II ChIP were performed in 50 ml canonical tube with 65 µg of anti-RNA Polymerase II 8WG16 monoclonal antibody (# MMS-126R, Covance). Immunoprecipitated chromatin was washed once with TSEI buffer, once with TSEII buffer, once with LiCl Buffer III, and once with TET buffer (see above for buffer composition). Beads were finally resuspended in 2 ml of 1X TE buffer supplemented with 1X protease inhibitor cocktail.
iii. A sample corresponding to 1/40^th^ of each ChIP was set apart and processed for DNA purification and assessment of ChIP enrichment by qPCR. Enrichment was calculated as the ratio between *Aldob* TSS ChIP signal normalized to input and intergenic region ChIP signal normalized to input. The *Aldob* primer sequences used are (forward) TGTTATCATTAACCCAGCTTGC and (reverse) CTGCCACCTCACACAGCTT. The intergenic region primer sequences are described above.

Libraries were generated following published protocols (50-52). Immunoprecipitated chromatin was end-repaired, processed for ligation with biotinylated half-linkers, and 5’ phosphorylated while still complexed with Pol II antibodies on magnetic beads. Chromatin was then eluted, diluted to a final volume of 10 ml, and used for the proximity ligation step (performed for a minimum of 16 hours at 4°C under extremely diluted conditions, i.e., < 0.2 ng DNA/µL, to favor ligation events within individual crosslinked chromatin complexes). Following the proximity ligation step, chromatin was treated with proteinase K, reverse crosslinked, and the DNA purified. ChIA-PET DNA was then digested with MmeI, immobilized on streptavidin beads, and ligated to the ChIA-PET adapters. The DNA samples were finally proceeded through nick translation and a PCR amplification step to generate the library (number of cycles indicated in Fig. S7A). Libraries were ran on an agarose gel, and fragment of ~229 bp were gel-extracted, purified, and quantified using a quantus fluorometer.

### Sequencing and computational analysis of Pol II ChIA-PET libraries

#### Sequencing and read alignment to the mouse genome

ChIA-PET libraries were sequenced on a HiSeq 2000 (JM08 and JM12) or a MiSeq (JM11) to a length of 90 bp (JM08), 100 bp (JM11), and 125 bp (JM12). Reads from the fastq files were processed to extract the tag 1 and 2 (i.e., read 1 and 2) along with their accompanying half-linker code using a custom-made Python script and generate two fastq files (R1 and R2 file) containing the sequence identifier, the raw sequence, and the sequence quality values for each tag. These two files were then aligned to the mouse genome (mm10 version) as paired-end reads using bowtie2 and the options -X 5 and --fast.

#### Paired-End Tags (PETs) filtering

Only paired tags with both reads mapping uniquely to the mouse genome were considered in our analysis. First, PETs were parsed based on the half-linker barcodes into non-chimeric PETs (specific products) or chimeric PETs (non-specific products) (Fig. 6A). Then, duplicated PETs (i.e., PCR duplicates) were removed for both chimeric and non-chimeric products. PETs with a tag location shifted by 1 bp compared to an existing PET were also considered as PCR duplicates and removed. Identical PET found at both ZT6 and ZT18 for the same experiment (JM08, JM11 or JM12), and which likely originate from PCR errors due to priming from half-linkers, were also filtered out. Quality of the ChIA-PET experiments was assessed by determining the percentage of non-chimeric/specific *vs*. chimeric/non-specific PETs (Fig. S7C), the proportion of PETs with both tags being on the same chromosome (Fig. S7D), and the distance between each tag (Fig. S7B, D).

#### Functional analysis of the mouse liver Paired-End Tags (PETs)

PETs with both reads on the same chromosome and with a distance between reads ≥ 500 bp were only considered in our analysis (n = 218,639 PETs, Table S4). For all PETs, each of the two tags was extended to 200 bp (tag location ± 100 bp) and this tag genomic location (chr:start-end) was used to map tags to (i) a gene, and (ii) a DNase I hypersensitive site, using intersectBed from bedtools (45). Gene coordinates were defined at TSS - 10kb to TTS + 1kb. Mapping to DHS was performed using a more stringent analysis of the mouse liver DNase-Seq datasets from ENCODE (Table S5), and which mostly reports stronger CREs (see Fig. 6D). This is because relaxation of the parameters for DNase-Seq analysis often consolidate several distinct DHS peaks into one DHS of several kilobases, thereby resulting into reporting PETs into the same DHS when both reads were distinctively located into two different DHS peaks. DHS harboring a mouse liver BMAL1 peak were identified with intersectBed between the DHS peak list described above and the list of mouse liver ChIP-Seq peaks generated in this manuscript (Table S1). To validate that mouse liver PETs contribute to gene transcription (Fig. 6B, 6C), we used public mouse liver Nascent-Seq datasets (29), and averaged values for each of the 12 independent samples. Finally, we considered genes to be rhythmically expressed based on the analysis of the microarray datasets from Zhang and collaborators (8), as described above.

#### Analysis of H3K27ac ChIP-Seq signal at mouse liver Paired-End Tags (PETs)

Quantile normalized (see above) and log2-transformed values for H3K27ac ChIP-Seq signal at the 67,433 mouse liver DHS peaks were parsed based on their respective group (rhythmic or arrhythmic target gene expression, DHS with or without a BMAL1 ChIP-Seq peak, *etc*), and averaged.

### Statistical analysis

Statistical analysis was carried out in JMP^®^, Version 12.0.1 SAS Institute., Cary, NC, 1989-2007. ChIP-Seq and DNase-Seq signals were analyzed using a Kruskal-Wallis test, and post-hoc analysis with a Wilcoxon each pair test. Analysis of TF mRNA expression between the three tissues was performed using a one-way ANOVA. Analysis of the differences in BMAL1 peaks genomic locations was performed using a chi-square test, and differences in the number of BMAL1 peaks per genomic locations we analyzed by a Fisher’s exact test. Spearman correlation was used to determine the degree of correlation between signals (e.g., ChIP-Seq with DNA-Seq) or signal between tissues. Results were considered significant if p-value < 0.05 for the Kruskal-Wallis and ANOVA tests, and p-value < 0.01 for Fisher’s exact test.

### Data availability

The sequencing datasets generated in this paper (BMAL1 ChIP-seq and Pol II ChIA-PET) have been deposited to GEO under the accession code GSE110604.

## Acknowledgements

We thank Christopher Bradfield for kindly providing the *Bmal1^-/-^* mouse, Craig Kaplan, Christine Merlin, and Aldrin Lugena for insightful suggestions at various stages of the project, Christine Merlin, Deborah Bell-Pedersen and Paul Hardin for comments on the manuscript, the Texas A&M Institute for Genome Sciences and Society for providing access and maintenance to their high-performance computing cluster, Michael Rosbash, Kate Abruzzi, and Ryanne Spann for helping with sequencing BMAL1 ChIP-Seq libraries, and Charles Johnson and Richard Metz for helping with sequencing the ChIA-PET libraries. We are also grateful to Paul Hardin and Matthew Sachs laboratories for technical support. This work was supported by startup funds from Texas A&M University. The laboratory of S.H.Y. is supported by the National Institute of General Medical Sciences (R01GM114424), and the laboratory of Z.C. by the Robert A. Welch Foundation (AU-1731) and the National Institute on Aging (R01AG045828). J.S.T. is an Investigator in the Howard Hughes Medical Institute.

## Contributions

J.R.B. and J.S.M. conceived and designed the research. J.R.B. performed most of the experiments and the bioinformatics analysis. J.S.M performed the ChIA-PET experiments with the help of A.J.T and J.R.B. B.G., C.A.O., and J.S.M. helped with the bioinformatics analysis. J.S. and H.V. performed chromatin immunoprecipitations at early stage of the project. S.H.Y., Z.C., and J.S.T. contributed to the BMAL1 antibody. N.G. helped with the ChIA-PET bioinformatics analysis. J.R.B. and J.S.M. wrote the manuscript.

## Supplementary Tables

Table S1: genomic location of all 9066 BMAL1 peaks and their target genes

Table S2: motif analysis at BMAL1 peaks (group 5, 6, 7, and group 1 separately).

Table S3: motif analysis at footprints. Table S3A: BMAL1 peak center +/- 100bp (group 5, 6, 7).

Table S3B: motif analysis at footprints within BMAL1 DHS (group 5, 6, 7)

Table S4: ChIA-PET analysis and all annotated PETs with reads on the same chromosome.

Table S5: genomic location of all 67,443 DHS peaks in the mouse liver used for the analysis of DHS interactions.

## Supplementary Figures

Figure S1-S7

**Supplementary Figure 1.**
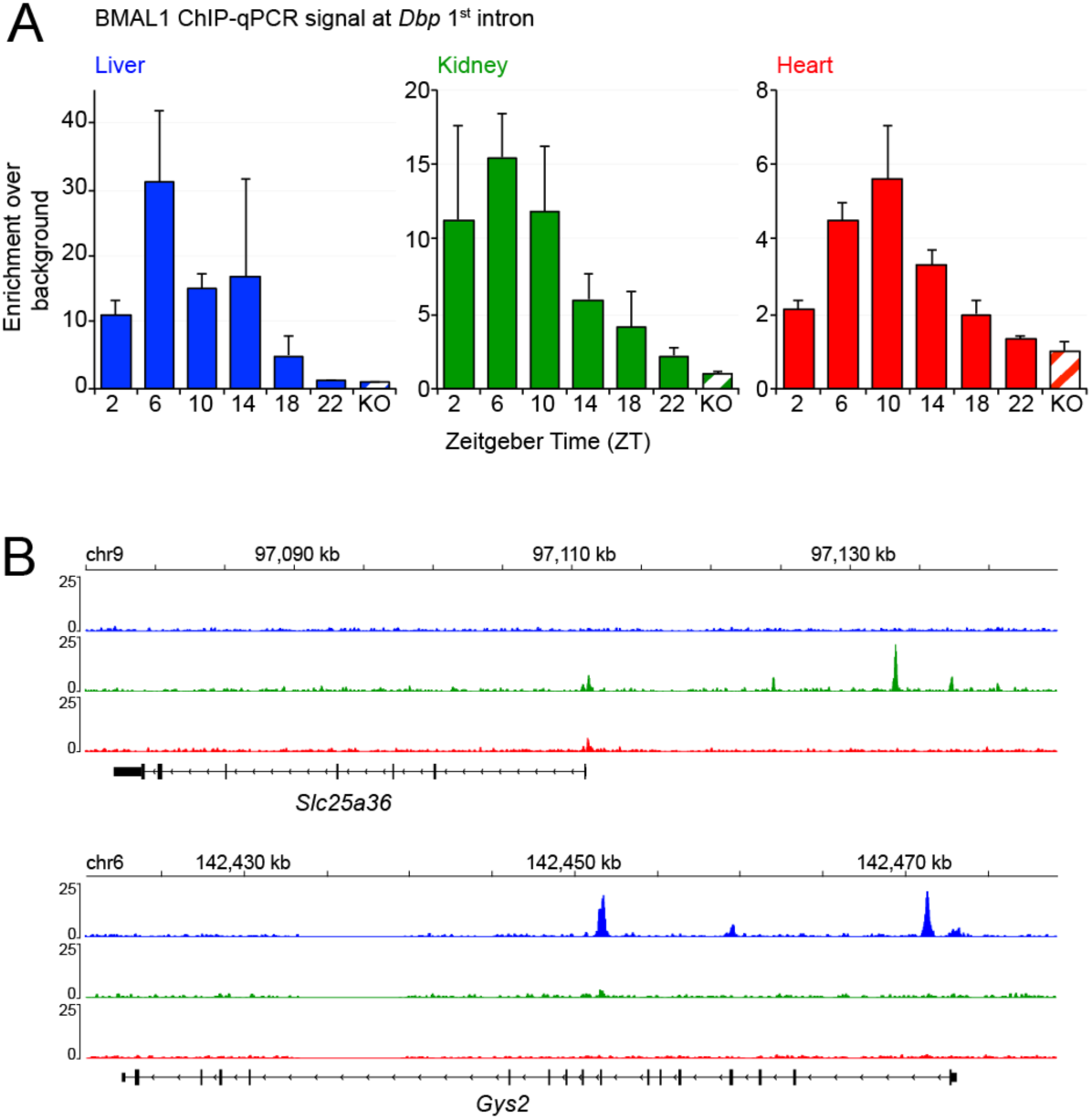
**BMAL1 cistromes are largely tissue specific**. (related to Fig. 1) **(A)** BMAL1 ChIP-qPCR signal at *Dbp* 1^st^ intron over the course of the day in the mouse liver (blue), kidney (green), and heart (red). Tissues were collected in wild-type mice at ZT2, ZT6, ZT10, ZT14, ZT18, and ZT22, and in *Bmal1* knockout mice at ZT6 for negative control. ChIP-qPCR signal normalized to the input signal corresponds to the average ± s.e.m. of 3 biological replicates, and the ratio ChIP/input was set to 1 for the *Bmal1* knockout mice samples. **(B)** Genome browser view of BMAL1 ChIP-Seq signal in the liver (blue), kidney (green), and heart (red) at *Slc35a6* and *Gys2* gene loci. BMAL1 ChIP-Seq signal is tissue-specific, with kidney-specific BMAL1 binding at *Slc35a6*, and liver-specific BMAL1 signal at *Gys2*.

**Supplementary Figure 2.**
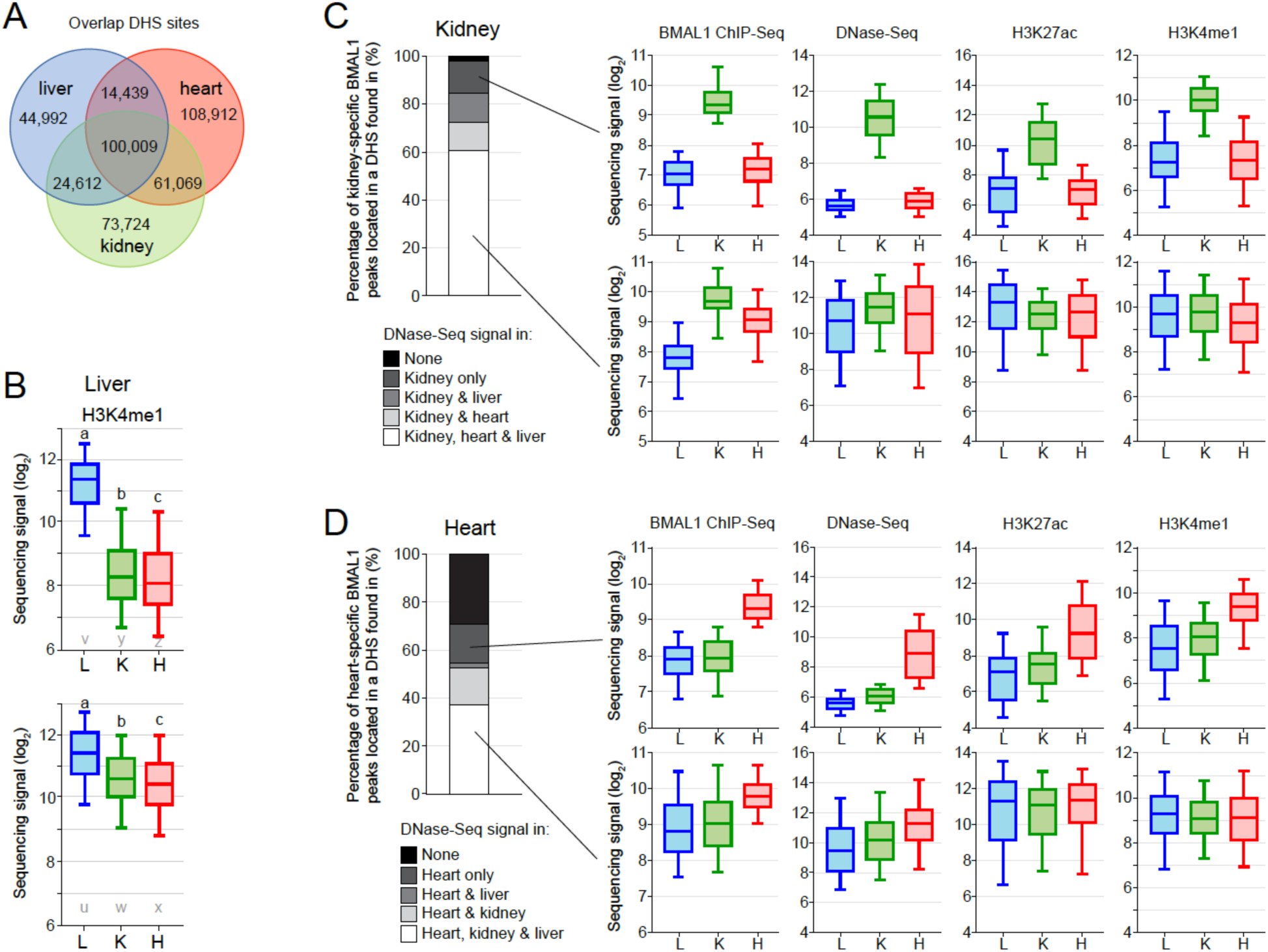
**The chromatin environment shapes tissue specific BMAL1 binding**. (related to Fig. 2) **(A)** Venn diagram depicting the overlap of DNase hypersensitive sites (DHS) between the mouse liver, kidney, and heart. DNase-Seq datasets were downloaded from the ENCODE project, and analyzed to define DHS in each of the three tissues. Of the total 427,748 DHS among the mouse liver, kidney, and heart, 100,009 are common to all three tissues. **(B)** H3K4me1 ChIP-seq signal (log2 scale) in the mouse liver, kidney, and heart at liver-specific BMAL1 peaks located in liver-specific DHS (top), or at liver-specific BMAL1 peaks located at DHS common to the liver, kidney and heart. H3H4me1 ChIP-Seq datasets were downloaded from the ENCODE project, and signal was calculated at BMAL1 peak center ± 1kb. **(C)** Left: Stacked bar chart representation of the percentage of kidney-specific BMAL1 peaks located within kidney-specific DHS or DHS found in other tissues. Right: Boxplot representation of BMAL1 ChIP-seq, DNase-seq, H3K27ac ChIP-seq and H3K4me1 ChIP-seq signals at kidney-specific BMAL1 peaks located at a kidney-specific DHS or at a DHS common to the liver, kidney and heart. Signal for BMAL1 ChIP-seq and DNase-seq was calculated at BMAL1 peak center ± 250 bp, and at BMAL1 peak center ± 500 bp for ChIP-seq of histone modifications. **(D)** Similar to panel C, but for heart-specific BMAL1 peaks.

**Supplementary Figure 3.**
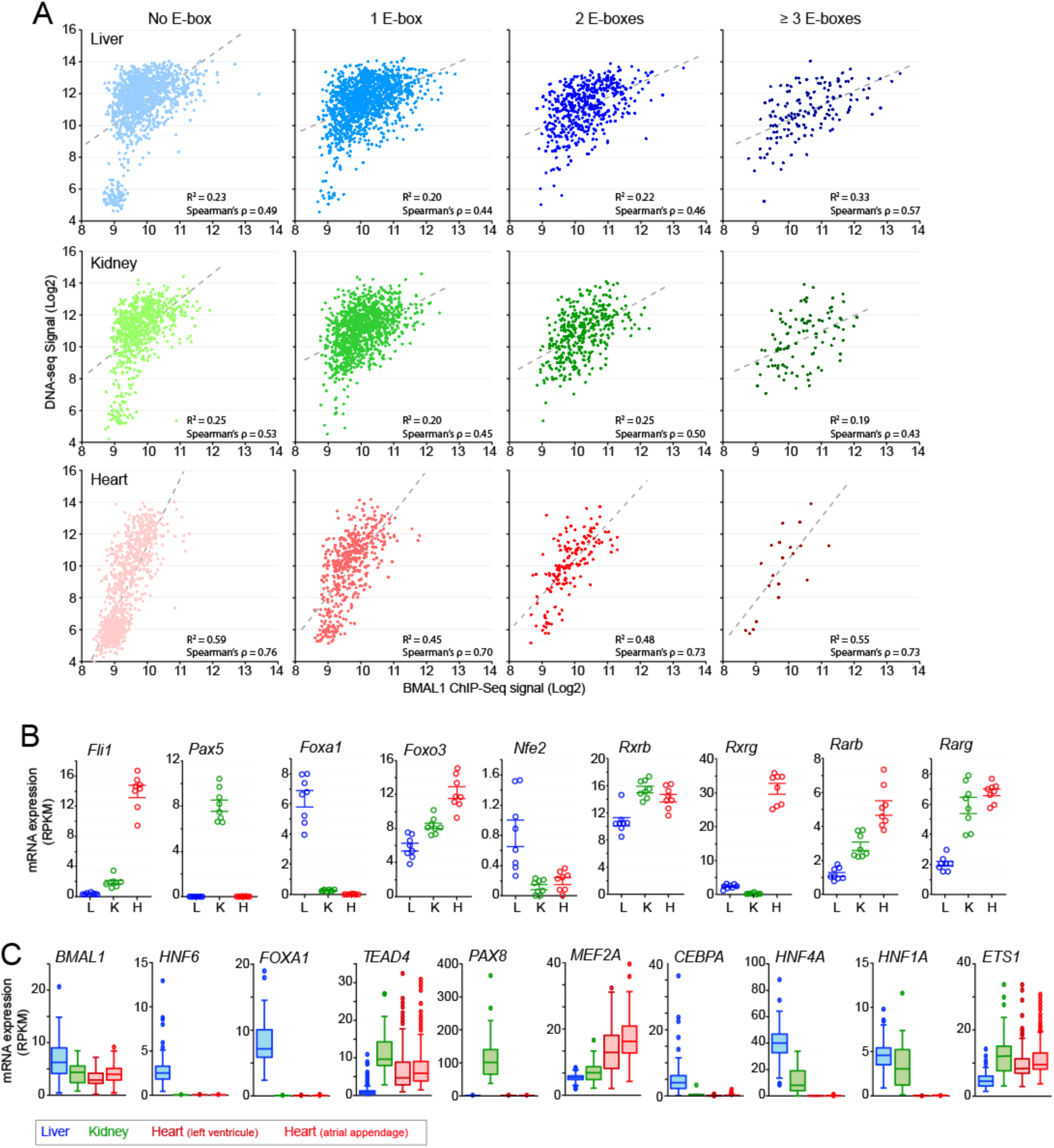
**Tissue-specific transcription factors may contribute to tissue-specific BMAL1 DNA binding**. (related to Fig. 3) **(A)** Correlation between DNase-seq signal and BMAL1 ChIP-seq signal for liver- (blue), kidney- (green) and heart- (red) specific BMAL1 peaks parsed based on the number of E-boxes. The E-boxes used in the analysis were CACGTG, CACGNG, and CACGTT. BMAL1 ChIP-seq signal and DNase-seq signal, which are represented in log2 scale, were calculated at BMAL1 peak center ± 250 bp. **(B)** mRNA expression in the mouse liver, kidney, and heart of *Bmal1* and transcription factors whose motifs were enriched at BMAL1 ChIP-Seq peaks. mRNA expression was calculated using public RNA-Seq datasets (8) from samples collected over the course of the 24-hr day. **(C)** mRNA expression (RPKM) in human liver (blue), kidney (green), and heart (red, left ventricle and atrial appendage) for the tissue-specific transcription factors displayed in Fig. 3C, D, and E. Data were retrieved from the GTEx portal (https://www.gtexportal.org/home/) (20).

**Supplementary Figure 4.**
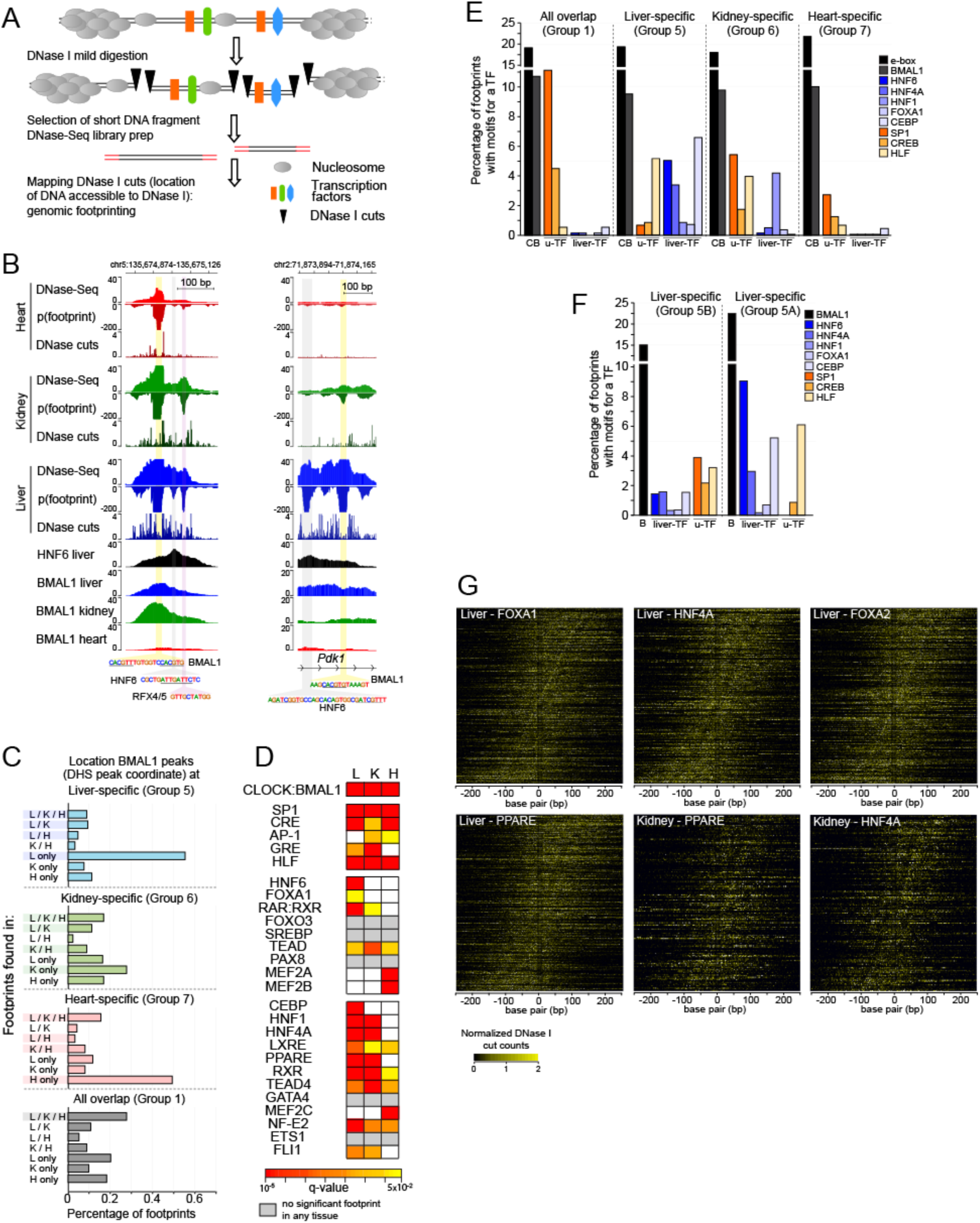
**DNase I footprint analysis indicates that tissue-specific transcription factors bind to tissue-specific BMAL1 DNA binding**. (related to Fig. 3) **(A)** Illustration of the DNase-Seq protocol, and the analysis of DNase I cuts that reveal footprints at DNase I hypersensitive sites (DHS). Mild DNase I digestion of nuclei preserves closed chromatin and transcription factors/nucleosomes-bound regions from DNase I cutting. Thus, mapping regions uncut by DNase I at DHS can reveal regions that are occupied by transcription factors and/or nucleosomes. **(B)** Genome browser view of DNase-seq signal, footprints p-value, DNase I cut sites, and BMAL1 ChIP-seq signal in the mouse liver (blue), kidney (green), and heart (red) at two BMAL1 DNA binding sites. DNase-Seq datasets were downloaded from the ENCODE project, and footprint p-value visualization files were generated using pyDNase (25, 48). Mouse liver HNF6 ChIP-Seq signal, which was downloaded from a public dataset (53), is also displayed. Both BMAL1 peaks exhibit significant footprints at motifs corresponding to E-boxes, as well as to motifs for other transcription factors including the liver-specific transcription factor HNF6. **(C)** Distribution of DNase I footprints detected at the genomic coordinate of the DHS bound by BMAL1, and parsed based on the tissue(s) they were found in. Analysis was performed at BMAL1 peaks that are liver-specific (blue), kidney-specific (green), heart-specific (red), or common to all three tissues (grey). **(D)** Motif enrichment of transcription factors performed at the DNase I footprints identified within the genomic coordinate of the DHS bound by BMAL1, for liver-, kidney-, and heart-specific BMAL1 peaks (footprint center ± 15 bp). Enrichments are displayed for the transcription factors shown in panel C if q-value < 0.05, and colored in grey in no significant footprint is detected in any of the three tissues. **(E, F)** Percentage of footprints for BMAL1 (black), liver-specific transcription factors (HNF6, HNF4A, HNF1, FOXA1, CEBP; blue), and ubiquitous transcription factors (SP1, CREB, HLF; orange) identified at (E) BMAL1 peaks common to all three tissues (group 1), BMAL1 peaks that are liver-specific (group 5), kidney-specific (group 6), and heart-specific (group 7), or at (F) liver-specific peaks that are located at a liver-specific DHS (group 5A) or located at a DHS that is common to the liver, kidney, and heart. **(G)** Heatmaps representing the DNase I cuts at BMAL1 peaks containing an E-box and a motif for another transcription factor. DNase I cut signal is centered on the E-box and sorted based on the distance between the E-box and the transcription factor motif. Scale corresponds to E-box center ± 250bp.

**Supplementary Figure 5.**
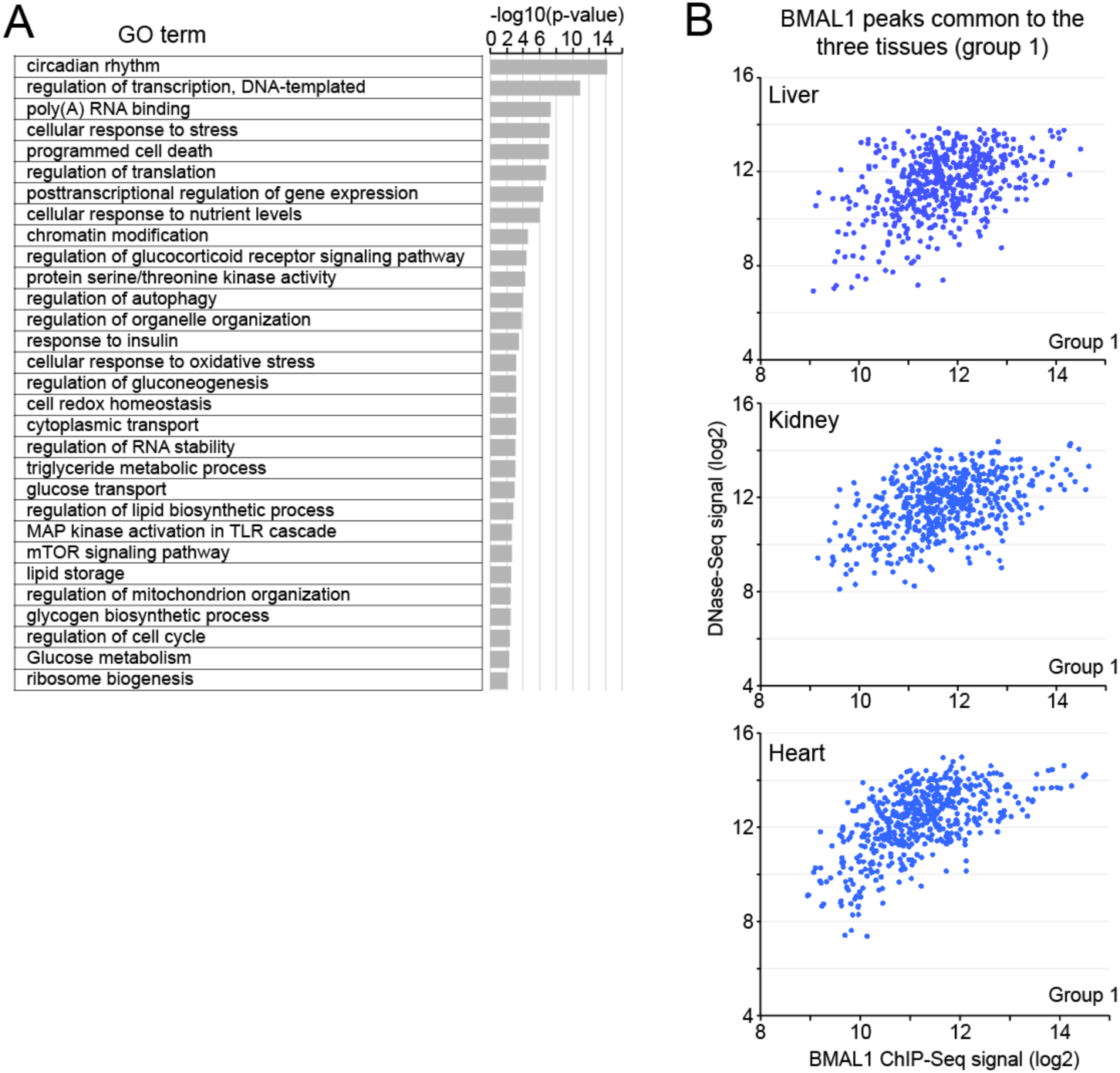
**BMAL1 peaks common to all three tissues target genes regulating general biological functions, and exhibit stronger ChIP-seq signal** (related to Fig. 4) **(A)** Gene ontology analysis of the genes targeted by BMAL1 peaks common to all three tissues (p-value < 0.05). **(B)** Correlation between DNase-seq and BMAL1 ChIP-seq signals for BMAL1 peaks that are common to all three tissues. Signal was calculated at BMAL1 peak center ± 250 bp in the mouse liver, kidney, and heart, and is displayed in log2 scale.

**Supplementary Figure 6:**
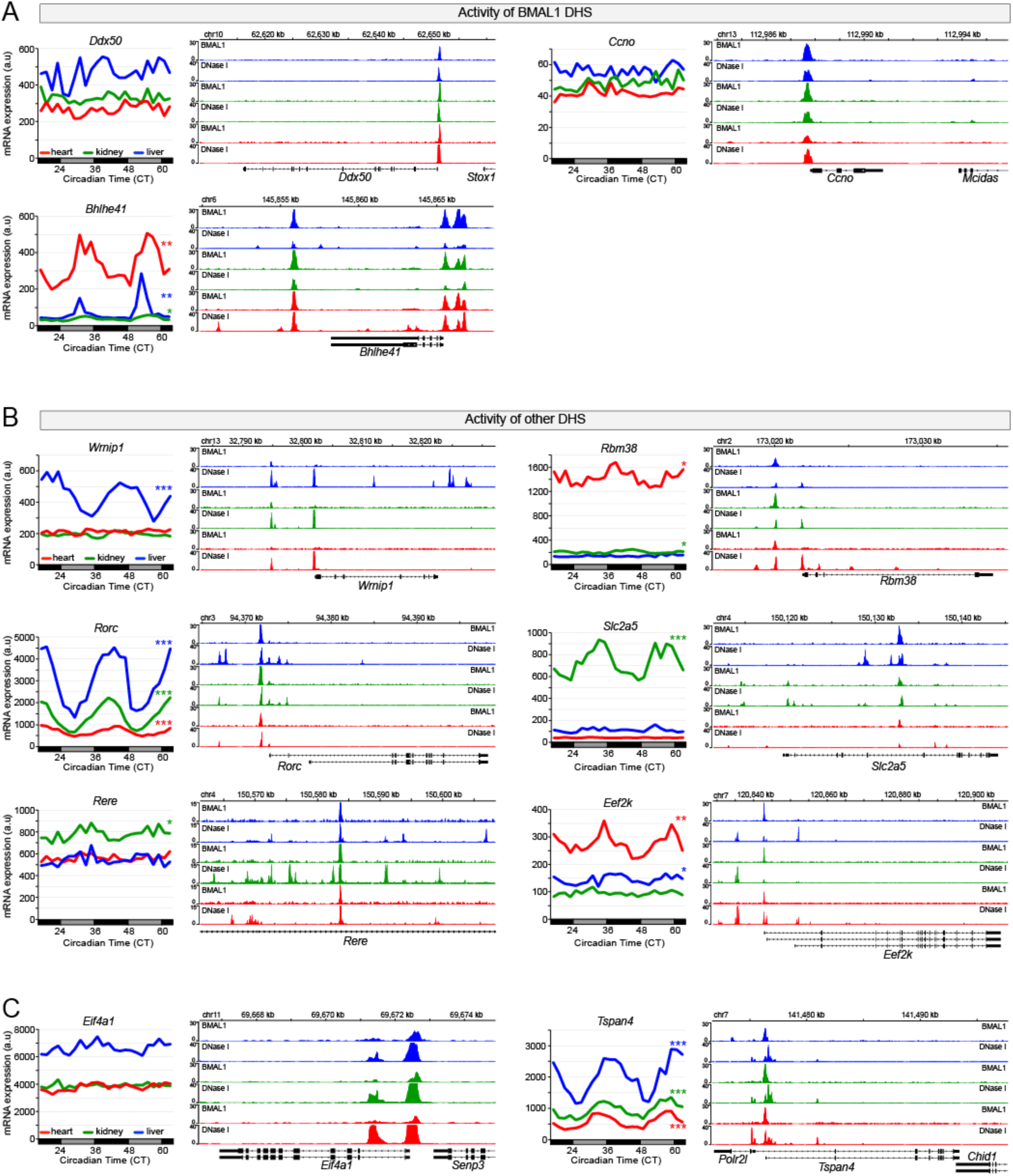
**Tissue-specificity of rhythmic BMAL1 target gene expression relies on the transcriptional activities of DHS bound by BMAL1, but also other DHS**. (related to Fig. 5) **(A-C)** mRNA expression (left) and genome browser view (right) of BMAL1 ChIP-seq and DNase-seq signals in the mouse liver (blue), kidney (green), and heart (red). Rhythmic expression determined by JTK-cycle is defined as *** if q-value < 0.001, ** if q-value < 0.01, and * if q-value < 0.05. Panel A represents examples for which the activity of BMAL1 DHS likely contributes to the differences in mRNA expression, whereas panels B and C represent examples for which the activity of other DHS likely contributes to BMAL1-mediated rhythmic transcription. The activity of these other enhancers often enhances (rhythmic) target gene expression (B), but it can also lead to decrease (rhythmic) gene expression (C).

**Supplementary Figure 7:**
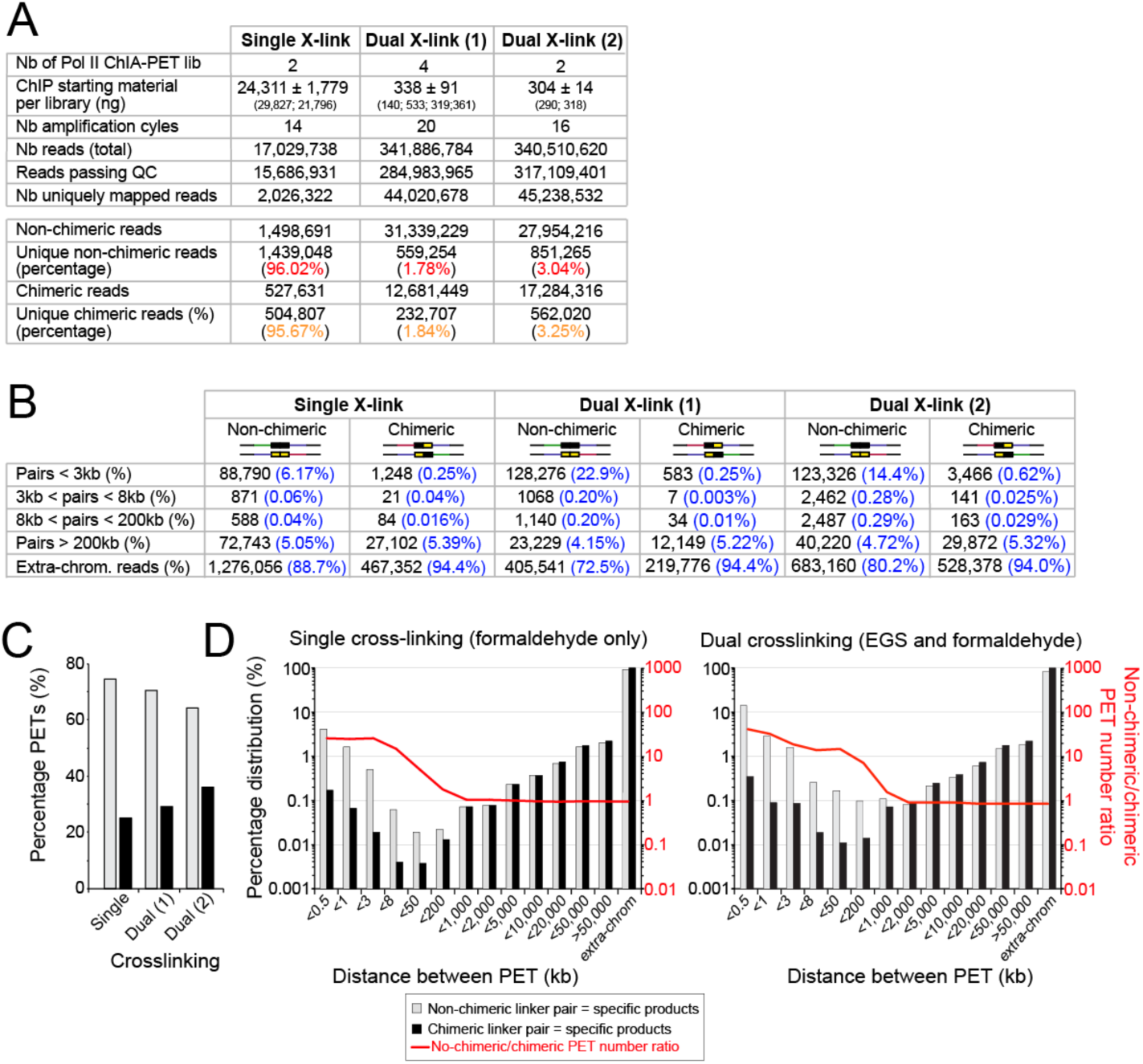
**Pol II ChIA-PET datasets uncover interaction between DHS in the mouse liver**. (related to Fig. 6) **(A)** Summary of the sequencing analysis for the three independent Pol II ChIA-PET experiments performed in this project. QC stands for quality check. **(B)** Number (black) and percentage (blue) of Paired-End Tags (PET) based on the distance between the two reads for the three independent Pol II ChIA-PET experiments. Reads mapping to two different chromosomes are labeled as extra-chrom. reads. **(C)** Percentage of PET with non-chimeric half-linker barcodes (specific products, grey) and chimeric half-linker barcodes (non-specific products, black) for each of the three independent Pol II ChIA-PET experiments. **(D)** Distribution of the PET length for the non-chimeric PET (specific products, grey) and the chimeric PET (non-specific products, black), for the mouse liver Pol II ChIA-PET experiments performed with single crosslinked nuclei (left) or dual crosslinked nuclei (right). The ratio between non-chimeric PET and chimeric PET is overlaid in red. Both y-axis are represented in log_10_ scale.

